# Reproducible and predictable reorganization of place fields driven by grid subfield rate changes

**DOI:** 10.64898/2026.02.10.705142

**Authors:** Christine M. Lykken, Benjamin R. Kanter, Jasmine Kaslow, Oscar M. T. Chadney, Kadjita Asumbisa, Lucie A. L. Descamps, Clifford G. Kentros

**Affiliations:** Kavli Institute for Systems Neuroscience and Centre for Algorithms in the Cortex, Norwegian University of Science and Technology, Olav Kyrres gate 9, 7030 Trondheim, Norway; Institute of Neuroscience, University of Oregon, 1254 University of Oregon, Eugene, Oregon 97403, USA

**Keywords:** hippocampus, medial entorhinal cortex, chemogenetics, place cells, grid cells

## Abstract

Understanding how the brain constructs stable yet flexible maps of space remains a central challenge in neuroscience. Place cells in the hippocampus fire at specific locations in a given environment, but reorganize completely upon introduction to another environment in a process called remapping. The medial entorhinal cortex (MEC) provides a major cortical input to the hippocampus, and the spatially periodic firing patterns of its grid cells are thought to contribute to place field formation. We previously showed that chemogenetic depolarization of MEC layer II stellate cells selectively altered firing rates within individual grid cell subfields, impaired spatial memory, and induced a form of reversible place cell remapping that we called artificial remapping. However, it remains unclear whether artificial remapping reflects a reproducible and stable mapping from entorhinal inputs to place cell outputs or a random reorganization of place fields. To explore the transfer of information between MEC and hippocampus, we repeated this chemogenetic manipulation on consecutive days and found that stimulating the same stellate cells produced similar changes in both grid subfield rates and place field locations. Using both experimental and simulated data, we show that baseline place cell activity patterns could be used to predict place field locations following the manipulation. These findings provide direct evidence for consistent input-output relationships in the entorhinal-hippocampal system and point to a central role for grid subfield rate changes in the reorganization of hippocampal spatial representations.

## Introduction

The hippocampus is essential for spatial navigation and episodic memory, or our ability to recall the ‘what’, ‘when’, and ‘where’ features of an experience (Scoville and Milner, 1957; Eichenbaum, 2017). Place cells in the hippocampus fire at specific locations within an environment and are thought to generate a cognitive map that guides behavior and supports navigation (O’Keefe and Dostrovsky, 1971; O’Keefe and Nadel, 1978; Eichenbaum et al., 1999; Robinson et al., 2020). When an animal enters a different environment, place cells remap, exhibiting unpredictable changes in firing rate and/or location (i.e., global remapping; Muller and Kubie, 1987; Bostock et al., 1991; Leutgeb et al., 2005). This orthogonalization is critical because it enables the hippocampus to store a large number of discrete representations with a limited number of cells, a necessary feature of a high-capacity episodic memory network (Treves and Rolls, 1991). Together, the formation of these precise spatial representations and their orthogonalization across contexts are thought to underlie hippocampal-dependent spatial memory (Kentros, 2006; Kanter et al., 2017; Robinson et al., 2020). However, it remains unclear exactly how specific patterns of input to the hippocampus give rise to stable yet flexible place cell representations.

The primary input to the hippocampus is the entorhinal cortex (Cappaert et al., 2015). Grid cells, which fire in a periodic hexagonal pattern that tiles the environment, are located in the superficial layers of the medial entorhinal cortex (MEC), along with several other spatially- and/or directionally-modulated cell types (Taube et al., 1990; Hafting et al., 2005; Sargolini et al., 2006; Savelli et al., 2008; Solstad et al., 2008; Diehl et al., 2017). Over the past two decades, both empirical and computational work have focused on the role of grid cells in the generation and remapping of place fields (Solstad et al., 2006; Fuhs and Touretzky, 2006; McNaughton et al., 2006; Rolls et al., 2006; Fyhn et al., 2007; Treves, 2009; de Almeida et al., 2009; Savelli and Knierim, 2010; Monaco and Abbott, 2011; Ormond and McNaughton, 2015; Kanter et al., 2017; Lykken et al., 2025). When place cells remap between distinct environments, grid cells belonging to different modules (i.e., groups of grid cells with similar spacing and orientation) undergo unique changes in spatial phase (Lykken et al., 2025). Varying degrees of remapping have also been observed in the absence of changes in grid phase (Kanter et al., 2017; Diehl et al., 2017; Lykken et al., 2025), following disruptions of grid cell activity (Brandon et al., 2014), and even after removal of the MEC (Schlesiger et al., 2018), suggesting that additional mechanisms may contribute to the positioning of place fields. For instance, chemogenetic depolarization of MEC layer II neurons (MEC LII) altered the firing rates within individual fields of grid cells (i.e., grid subfields) and induced robust remapping of place cells, even when grid phase remained stable (Kanter et al., 2017). Similarly, computational models incorporating variability in grid subfield rates (rather than assuming uniform firing across subfields) have demonstrated that changing grid subfield rates can drive hippocampal remapping (Lyttle et al., 2013; Dunn et al., 2017). These observations raise the question of whether the transformation of entorhinal input to place cell output operates according to stable, predictable rules. In other words, does a particular change in entorhinal input consistently produce the same change in hippocampal output?

## Results

To address these questions, we used a transgenic mouse line that enables selective chemogenetic activation of excitatory neurons in MEC LII. Specifically, we crossed an hM3Dq-tetO DREADD (Designer Receptor Exclusively Activated by a Designer Drug) (Alexander et al., 2009) line to an Ent-tTA driver line (Yasuda and Mayford, 2006), which drives expression almost exclusively in reelin-positive stellate cells of MEC LII, targeting approximately 27% of this population (Kanter et al., 2017) (Figure S1). Double-positive offspring are hereafter referred to as hM3 mice. To assess how this manipulation altered entorhinal input and hippocampal output, naïve adult mice were implanted with chronic tetrode arrays targeting superficial MEC or dorsal CA1 (Figure S2). We recorded neural activity in these regions as mice freely explored a familiar open-field environment before and after intraperitoneal injection of the designer ligand clozapine-N-oxide (CNO) on two consecutive days.

### Chemogenetic activation of MEC LII elicits reproducible changes in MEC activity across days

Each recording day consisted of a thirty-minute baseline (BL) session followed by a two-hour session immediately after an injection of CNO (1 mg/kg) or saline in hM3 mice or littermate controls. As reported previously (Kanter et al., 2017), chemogenetic depolarization of MEC LII neurons in hM3 mice significantly altered the firing rates and field sizes of putative excitatory neurons relative to controls on both days (rate change, hM3 vs. Con: BL1×CNO1, hM3 n= 61, Con n = 46, Z = 4.8, p = 8.8 × 10^-7^; BL2×CNO2, hM3 n = 58, Con n = 39, Z = 5.0, p = 3.3 × 10^-7^; size change, hM3 vs. Con: BL1×CNO1, hM3 n = 46, Con n = 42, Z = 1.7, p = 0.04; BL2×CNO2, hM3 n = 41, Con n = 33, Z = 2.5, p = 6.6 × 10^-3^; one-sided Wilcoxon rank sum tests; Figures 1A and S3A-B). Here, we observed that the magnitude of these effects was highly correlated across days (rate change, BL1×CNO1 vs. BL2×CNO2: n = 49, r = 0.71, p = 1.0 × 10^-8^; size change, BL1×CNO1 vs. BL2×CNO2: n = 30, r = 0.76, p = 9.8 × 10^-7^; linear correlations; Figures 1A and S3C-D), indicating that our manipulation produced consistent changes in MEC activity. Importantly, the spatial firing patterns of MEC neurons, including grid cells, remained stable following CNO injection (spatial correlation, hM3 vs. Con: BL1×CNO1, hM3 n = 48, Con n = 42, Z = 1.6; p = 0.10; BL2×CNO2, hM3 n = 43, Con n = 34, Z = 1.8; p = 0.07; two-sided Wilcoxon rank sum tests; Figures 1A and S3E), confirming that these rate and field size changes occurred without dramatically reshaping the spatial input to the hippocampus.

**Figure 1.**
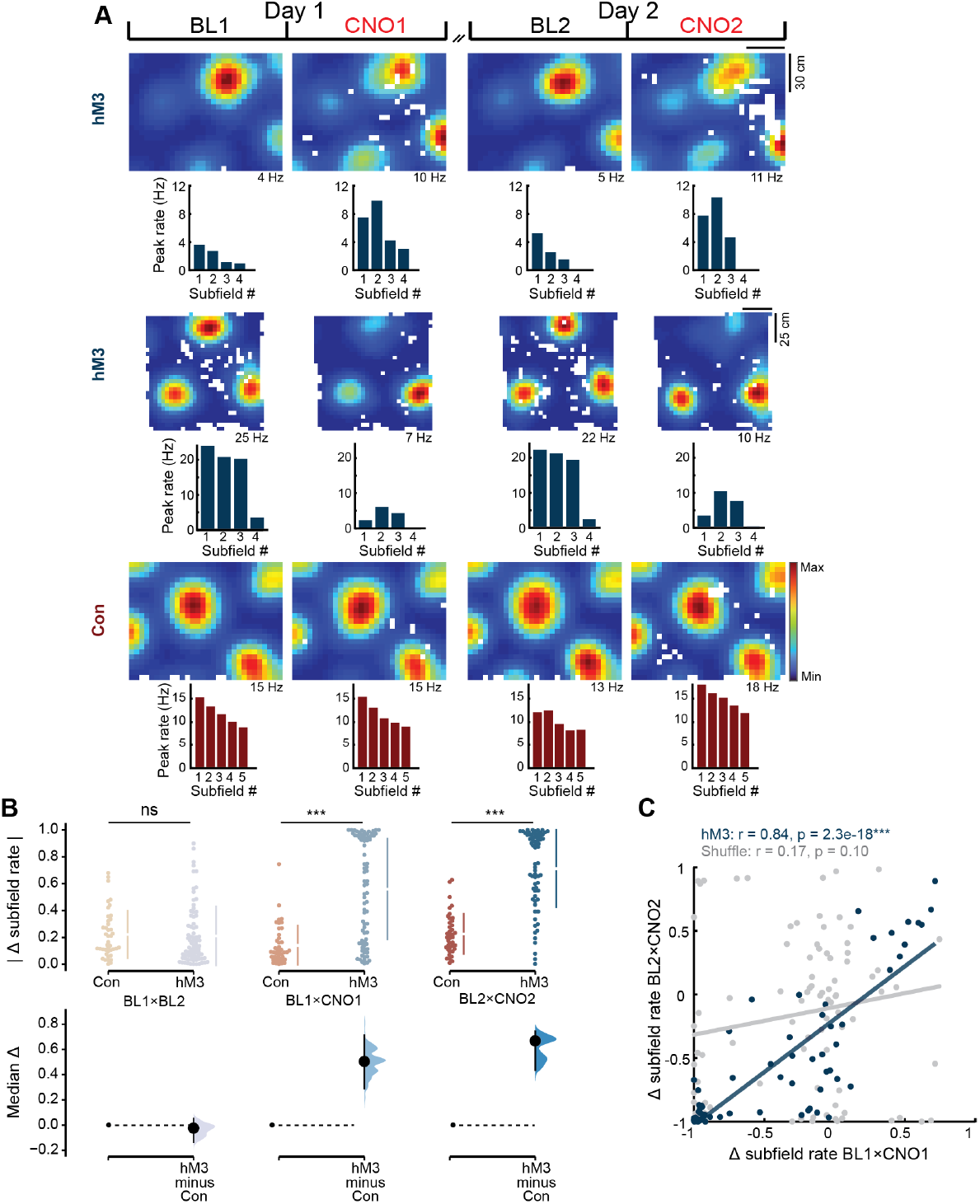
Depolarization of MEC LII on consecutive days produces consistent changes in grid subfield rates. **A)** Firing rate maps of representative grid cells in hM3 mice (top two rows) and a control mouse (Con, bottom row) before and after CNO injection on consecutive days. Color indicates firing rate. Peak firing rate is noted below each rate map. Peak firing rate within each grid subfield is shown below the rate map for that session. BL1, baseline session on Day 1; CNO1, 30-60 min post-CNO injection on Day 1; BL2, baseline session on Day 2; CNO2, 30-60 min post-CNO injection on Day 2. Dashed black lines indicate that BL2 was initiated 12+ hrs. after CNO injection. **B)** Panel shows grid subfield rate changes between sessions in hM3 (blue) and Con (red) mice. Left, there was no difference between hM3 and Con mice in grid subfield rate changes between BL sessions (hM3 vs. Con: BL1×BL2, hM3 n = 80, Con n = 42, Z = 1.1, p = 0.27; two-sided Wilcoxon rank sum test). Middle, right, grid subfield firing rates changed significantly between BL and CNO sessions on both recording days in hM3 mice versus controls (hM3 vs. Con: BL1×CNO1, hM3 n = 79, Con n = 42, Z = 5.7, p = 5.2 × 10^-9^; BL2×CNO2, hM3 n = 67, Con n = 40, Z = 7.0, p = 1.5 × 10^-12^; one-sided Wilcoxon rank sum tests). Top, points represent individual subfields; gapped lines represent mean ± standard deviation. Change refers to an absolute difference score (see **Methods**). Bottom, black dot: median; black bars: 95% confidence interval; filled curve: sampling-error distribution. **C)** Scatterplot showing significant correlation between grid subfield rate changes between BL and CNO sessions on Day 1 and Day 2 in hM3 mice (blue, n = 53, r = 0.84, p = 2.3 × 10^-18^, linear correlation), but not in a shuffled control group (gray, n = 97, r = 0.17, p = 0.10, linear correlation). Points represent individual subfields. Change refers to a difference score. *\*\*\*p* < 0.001.

Given the prominence of grid cells in theoretical models and our prior finding that chemogenetic depolarization of MEC LII selectively alters grid subfield rates without changing the spatial or directional tuning properties of other entorhinal neurons (Kanter et al., 2017), we focused our subsequent analyses on grid cells. Among recorded grid cells, grid subfield firing rates changed significantly on both recording days in hM3 mice relative to controls (hM3 vs. Con: BL1×CNO1, hM3 n = 79, Con n = 42, Z = 5.7, p = 5.2 × 10^-9^; BL2×CNO2, hM3 n = 67, Con n = 40, Z = 7.0, p = 1.5 × 10^-12^; one-sided Wilcoxon rank sum tests; Figure 1B, middle and right), but were stable across baseline sessions in both groups (hM3 vs. Con: BL1×BL2, hM3 n = 80, Con n = 42, Z = 1.1, p = 0.27; two-sided Wilcoxon rank sum test; Figure 1B, left). These subfield rate changes were strongly correlated across days in hM3 mice, but not in control mice or a shuffled control dataset (see **Methods**; hM3 n = 53, r = 0.84, p = 2.3 × 10^-18^; Con n = 40, r = −0.05, p = 0.77; shuffle n = 97, r = 0.17, p = 0.10; linear correlations; Figures 1A and 1C). Rather than simply rescaling the magnitude of subfield firing rates, our manipulation also disrupted grid subfield rate relationships (i.e., the ordering of subfield rates from highest to lowest) on both days (Figure 1A). The alteration in subfield rate relationships was consistent across days, resulting in a significant correlation between grid field relationships during CNO1 and CNO2 in hM3 mice (CNO1 vs. CNO2, n = 66, r = 0.60, p = 2.0 × 10^-7^, linear correlation; Figures 1A and S3F), but not in a shuffled control group (see **Methods**; CNO1 vs. CNO2, n = 81, r = 0.03, p = 0.78, linear correlation). In control mice, subfield rate relationships were also stable over time (CNO1 vs. CNO2, n = 64, r = 0.57, p = 1.4 × 10^-5^, linear correlation; Figures 1A and S3F). Together, these results demonstrate that our chemogenetic manipulation reliably altered the subfield firing rates of grid cells in a similar manner across days without affecting their spatial phase, providing a means to systematically examine how this selective modulation of entorhinal inputs shapes downstream place field representations.

### Repeated depolarization of MEC LII produces consistent changes in CA1 activity

To determine whether consistent changes in entorhinal input were reflected in consistent changes in hippocampal output, we recorded place cells in dorsal CA1 while repeating the chemogenetic manipulation on two consecutive days. As in our MEC recordings, depolarizing MEC LII neurons in hM3 mice significantly altered the firing rates and field sizes of CA1 place cells relative to controls (rate change, hM3 vs. Con: BL1×CNO1, hM3 n = 113, Con n = 109, Z = 4.1, p = 1.8 × 10^-5^; BL2×CNO2, hM3 n = 109, Con n = 111, Z = 6.3, p = 1.6 × 10^-10^; size change, hM3 vs. Con: BL1×CNO1, hM3 n = 86, Con n = 101, Z = 6.1; p = 4.6 × 10^-10^; BL2×CNO2, hM3 n = 80, Con n = 102, Z = 5.5, p = 2.0 × 10^-8^; one-sided Wilcoxon rank sum tests; Figures 2A and S4A-B). The magnitude of these effects was highly correlated across days (rate change: n = 104, r = 0.60, p = 2.1 × 10^-11^; size change: n = 64, r = 0.45, p = 1.8 × 10^-4^; linear correlations; Figures 2A and S4C-D), indicating that chemogenetic manipulation of MEC LII consistently altered the firing properties of CA1 place cells across sessions.

**Figure 2.**
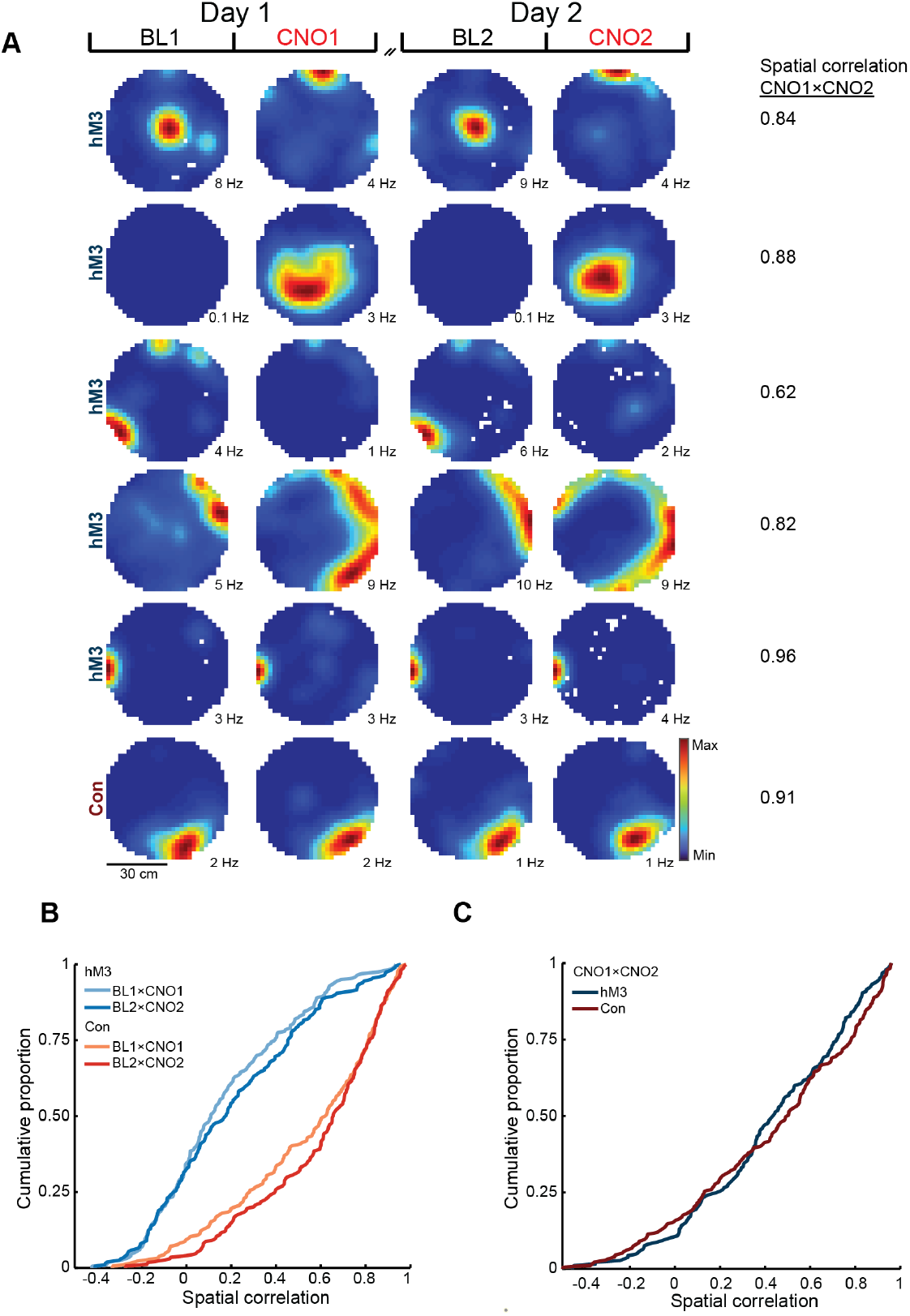
Depolarization of MEC LII on consecutive days produces a consistent reorganization of CA1 place cell activity. **A)** Firing rate maps of representative place cells in hM3 mice (top five rows) and a control mouse (Con, bottom row) before and after CNO injection on Day 1 and Day 2. Color indicates firing rate. Peak firing rate is noted below each rate map. Spatial correlation between rate maps from CNO sessions on each day is noted on the right for each cell. **B)** Cumulative distribution function shows the spatial correlation between place cell rate maps from BL and CNO sessions on each day in hM3 (BL1×CNO1, light blue; BL2×CNO2, medium blue) and Con (BL1×CNO1, light red; BL2×CNO2, medium red) mice. There was a significant decrease in the spatial correlation of place cells in hM3 versus Con mice on both days (hM3 vs. Con: BL1×CNO1, hM3 n = 106, Con n = 107, D* = 0.60, p = 1.6 × 10^-17^; BL2×CNO2, hM3 n = 108, Con n = 105, D* = 0.59, p = 7.5 × 10^-17^; two-sided Kolmogorov-Smirnov tests). **C)** Cumulative distribution function shows the spatial correlation between place cell rate maps from CNO sessions on each day in hM3 (dark blue) and Con (dark red) mice. There was no difference between hM3 and Con mice in the spatial correlation of place cells between CNO sessions (hM3 vs. Con: CNO1×CNO2, hM3 n = 97, Con n = 104, D* = 0.16, p = 0.15, two-sided Kolmogorov-Smirnov test), indicating that the reorganization of hippocampal place cell activity was consistent across days.

In addition, chemogenetically activating MEC LII neurons dramatically reshaped the spatial firing patterns of CA1 place cells on each recording day (Figure 2A). Spatial correlations between place cell rate maps from BL and CNO sessions on both days were significantly reduced in hM3 mice compared to controls (hM3 vs. Con: BL1×CNO1, hM3 n = 106, Con n = 107, D* = 0.60, p = 1.6 × 10^-17^; BL2×CNO2, hM3 n = 108, Con n = 105, D* = 0.59, p = 7.5 × 10^-17^; two-sample Kolmogorov-Smirnov tests; Figure 2B), consistent with a robust remapping of CA1 place cells on both days. In contrast, spatial correlations between rate maps from the BL session on each day did not differ between groups (hM3 vs. Con: BL1×BL2, hM3 n = 126, Con n = 114, D* = 0.10, p = 0.57, two-sample Kolmogorov-Smirnov test; Figure S4E), indicating that the hippocampal response to our manipulation was reversible. Crucially, there was also no difference between groups in spatial correlations between rate maps from the CNO session on each day (hM3 vs. Con: CNO1×CNO2, hM3 n = 97, Con n = 104, D* = 0.16, p = 0.15; two-sample Kolmogorov-Smirnov test; Figure 2C), indicating that repeated activation of MEC LII led to a consistent reorganization of hippocampal spatial representations. Taken together, these results demonstrate that consistent changes in entorhinal input drive reproducible transformations of hippocampal activity patterns, indicating that the entorhinal-hippocampal circuit transforms a given pattern of input according to stable and predictable rules.

### Predictable reorganization of place fields following depolarization of MEC LII

The reproducible nature of this transformation raises the possibility that particular changes in entorhinal input may bias place fields toward a predetermined subset of locations. It is still debated whether place cells receive broadly distributed synaptic inputs that allow them to form anywhere in the environment (Bittner et al., 2015, 2017; Diamantaki et al., 2018; Liao et al., 2024) or whether their inputs are biased toward particular locations (Lee et al., 2012; Cohen et al., 2017), forming a predetermined landscape that constrains where place fields can emerge. Although we do not measure changes in synaptic input directly, we hypothesized that if place cells receive biased rather than evenly distributed inputs, our manipulation might cause place fields to emerge in locations with pre-existing, but weak, firing activity. We therefore asked whether a place cell’s baseline activity pattern could be used to predict how its place field reorganizes following depolarization of MEC LII.

To test this, we examined each place cell’s baseline rate map and identified locations outside of the primary field that could serve as predictions of place field location following CNO injection (see **Methods**; Figures 3A and S5A). These predictions were generated by detecting local maxima in the baseline rate map and selecting up to three secondary peaks outside the primary field (see **Methods**). We primarily focused our analysis on place cells with fields that shifted between sessions (204/396 cells, 51.5%; criteria described in **Methods**). As shown previously (Ormond et al., 2023), place cells that were stable between sessions exhibited significantly higher baseline firing rates and larger field sizes than place cells that shifted between sessions (peak firing rate: stable vs. shift, D* = 0.19, p = 1.4 × 10^-3^; field size: stable vs. shift, D* = 0.16, p = 8.0 × 10^-3^; two-sided Kolmogorov-Smirnov tests), supporting the idea that stronger preexisting inputs may contribute to place field stability.

**Figure 3.**
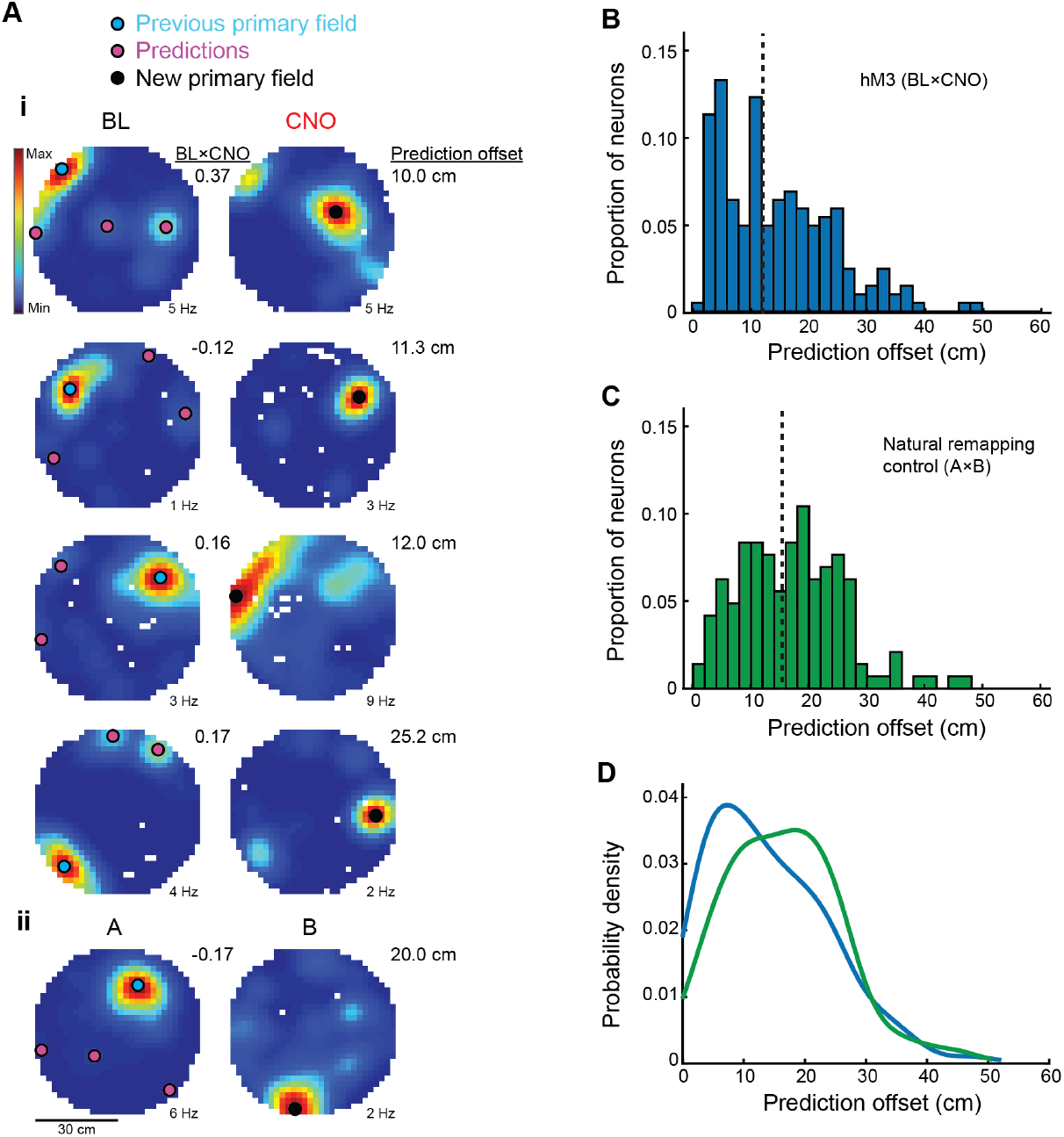
Predictable reorganization of CA1 place cell activity following depolarization of MEC LII. **Ai)** Firing rate maps of representative CA1 place cells in hM3 mice before and after CNO injection (one cell per row). Color indicates firing rate. Peak firing rate is noted below each rate map. For each cell, if the location of the primary place field shifted between BL (left, blue circles) and CNO (right, black circles) by more than 20 cm, we identified up to three secondary peaks outside of the primary field in the BL session (left, pink circles) that served as predictions of place field location. We then calculated the distance between each predicted location and the location of the primary place field in the CNO session, defining the shortest of these distances as the ‘prediction offset’. The spatial correlation between rate maps from the BL and CNO session for each cell is noted between columns. The prediction offset for each cell is shown on the right. Note that even when the spatial correlation between sessions was very low, we were often able to predict the location of the new place field for many place cells. **Aii)** Firing rate map of a representative CA1 place cell in a control mouse exposed to two distinct environments (A and B). Same convention as in Figure 3Ai. **B)** Histogram shows prediction offsets for all CA1 place cells recorded in hM3 mice (n = 204, median = 12.1 cm, 95% CI, 10.2 – 14.6 cm). **C)** Histogram shows prediction offsets for CA1 place cells recorded in a separate cohort of mice exposed to distinct environments (A×B, n = 144, median = 16.3 cm, 95% CI, 14.1 – 18.4 cm). Note that prediction offsets were significantly lower for place cells in hM3 mice than for place cells recorded as mice were moved between distinct rooms (hM3 vs. A×B, Z = 2.7, p = 3.4 × 10^-3^, one-sided Wilcoxon rank sum test), reflecting the predictable reorganization of place fields that occurs during artificial remapping. Vertical lines represent the median of each distribution. **D)** Kernel smoothed density estimate of prediction offset for place cells in hM3 mice (blue) and a separate cohort of mice exposed to distinct environments (green).

For those cells with fields that shifted between sessions, we then calculated the distance between each of the predicted locations and the cell’s primary field in the CNO session, defining the shortest of these distances as the ‘prediction offset’. As a control, we generated a shuffled dataset in which the predicted locations from each cell in the BL session were compared to the observed place field locations of randomly selected cells in the CNO session (see **Methods**). The median prediction offset was significantly lower for place cells in hM3 mice than for the shuffled dataset (hM3 n = 204, median = 12.1 cm, 95% CI, 10.2 – 14.6 cm; shuffle n = 261, median = 16.1 cm, 95% CI, 14.4 – 17.9 cm; hM3 vs. shuffle, Z = 3.9, p = 5.6 × 10^-5^, one-sided Wilcoxon rank sum test; Figures 3B and S5B). Prediction offsets were also significantly lower for hM3 mice than for a separate cohort of mice moved between two distinct environments (i.e. “natural” remapping, A×B n = 144, median = 16.3 cm, 95% CI, 14.1 – 18.4 cm; hM3 vs. A×B, Z = 2.7, p = 3.4 × 10^-3^, one-sided Wilcoxon rank sum test; Figures 3A and 3C-D), suggesting that artificial remapping differs from the seemingly random reallocation of place field locations observed during global remapping. We then confirmed that accurate predictions of place field location were possible even when place cells underwent extensive changes in firing rate and location. First, we determined that prediction accuracy was not simply driven by minimal changes in firing rate between sessions. Instead, we observed a clear reorganization of spiking activity from the original place field location in the BL session to the new place field in the CNO session (Figures S5C-D). Specifically, between sessions, there was a significant decrease in firing rate within the original place field (rate change within BL field: Con n = 320, median = −0.10; hM3 n = 204, median = −0.42; Con vs. hM3, Z = 8.4, p = 5.3 × 10^-17^, two-sided Wilcoxon sign-rank test; Figure S5C) and a corresponding increase within the new place field (rate change within CNO field: Con n = 320, median = 0.03; hM3 n = 204, median = 0.39; Con vs. hM3: Z = 8.4, p = 4.25 × 10^-17^, two-sided Wilcoxon sign-rank test; Figure S5D) in hM3 mice relative to controls. Moreover, there was no correlation between the firing rate in the predicted location in the BL session and the prediction offset (n = 204, r = −0.11, p = 0.12, linear correlation; Figure S5E), indicating that accurate predictions were possible even when the firing rate in the predicted location was very low. In line with this, spiking activity at the predicted location satisfied our criteria for place field detection for just 21.6% of place cells (44/204), consistent with the idea that our manipulation unmasked preexisting, subthreshold input. Second, we found that there was only a weak relationship between the prediction offset and the spatial correlation across sessions (n = 204, r = −0.26, p = 1.9 × 10^-4^, linear correlation; Figure S5F), indicating that successful predictions were possible even when the spatial correlation between sessions was very low. Therefore, despite extensive remapping, place field locations in CA1 could be reliably predicted from baseline activity patterns. Together, these results raise the possibility that predictable hippocampal reorganization arises from changes in input that reveal preexisting subthreshold place fields.

### Grid subfield rate changes drive predictable reorganization of place fields

Depolarization of MEC LII reconfigures the spatial input to the hippocampus by selectively altering the firing rates of individual grid cell subfields without changing their spatial phase (Figures 1A-B; Kanter et al., 2017). We therefore hypothesized that these rate-based modifications of grid subfields could drive the predictable reorganization of CA1 place fields observed here during artificial remapping. To test this idea, we incorporated our empirically observed grid subfield rate changes into a model of the grid-to-place cell transformation (de Almeida et al., 2012). An advantage of this model is that it does not incorporate reciprocal connections from place to grid cells, allowing us to isolate the influence of grid subfield rate changes on downstream place cell output.

Using a library of 10,000 simulated grid cells with variable subfield rates (see **Methods**), we first generated a set of simulated place cells with realistic place fields. The rate maps of this set of simulated grid and place cells represented the BL session (Figures 4A-B). We then modified the rate of each grid subfield by an amount drawn randomly from the distribution of subfield rate changes in hM3 mice before generating a second set of place cell rate maps (CNO session; Figures 4A-B and S6A). The peak firing rate of each simulated grid cell was held constant across sessions to isolate the effect of subfield rate changes on place field location.

**Figure 4.**
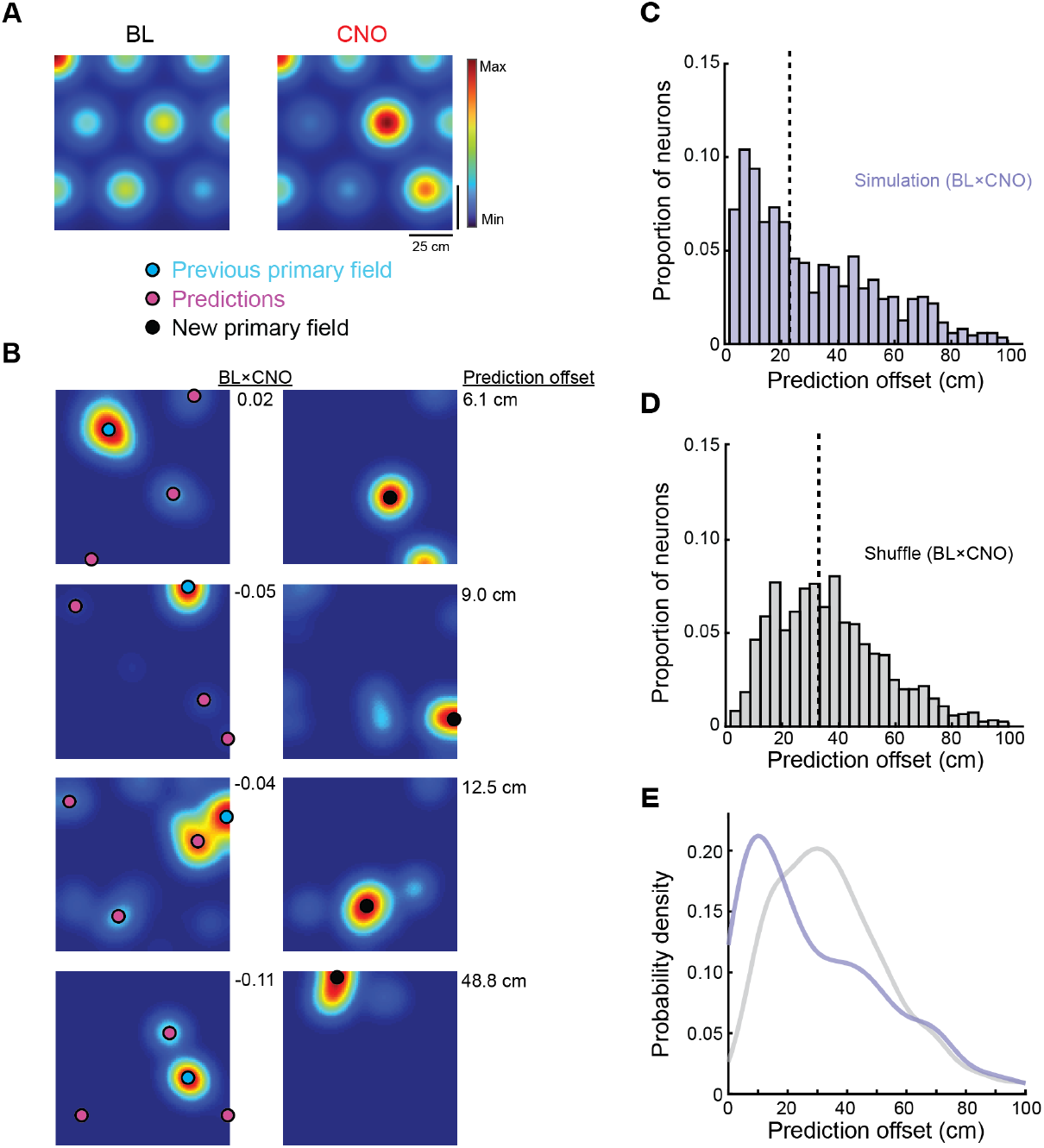
Grid subfield rate changes are sufficient to produce a predictable reorganization of hippocampal place fields. **A)** Firing rate maps of one simulated grid cell before and after modification of grid subfield rates. Color indicates firing rate. We first generated a library of 10,000 grid cells with variable subfield rates (left, BL). We then modified the rate of each subfield by an amount drawn randomly from a distribution of subfield rate changes in hM3 mice (right, CNO; see Figure S6A). The peak firing rate of each simulated grid cell was held constant across sessions to isolate the effect of subfield rate changes on place field location. **B)** Firing rate maps of four simulated place cells before (left, BL) and after (right, CNO) modification of grid subfield rates. Color indicates firing rate. Same convention for place field prediction as in Figure 3A. The spatial correlation between rate maps from the BL and CNO session for each cell is noted between columns. The prediction offset for each cell is shown on the right. **C-D)** Histograms show prediction offsets for simulated place cells (C, purple) and a shuffled control dataset (D, gray). Note that prediction offsets were significantly lower for simulated place cells than for the shuffled dataset (simulation n = 875, median = 23.0, 95% CI, 20.2 – 26.0 cm; shuffle n = 1,204, median = 33.0, 95% CI, 31.6 – 34.0 cm; simulation vs. shuffle, Z = 7.9, p = 1.3 × 10^-15^, one-sided Wilcoxon rank sum test). Vertical lines represent the median of each distribution. **E)** Kernel smoothed density estimate of prediction offset for simulated place cells (purple) and a shuffled control dataset (gray).

This manipulation of grid subfield rates produced a degree of remapping that closely matched our experimental observations (spatial correlation, BL×CNO: hM3 n = 394, median = 0.14, 95% CI, 0.09 – 0.17; simulation n = 1,975, median = 0.13, 95% CI, 0.11 – 0.16; hM3 vs. simulation, Z = 1.7, p = 0.09, two-sided Wilcoxon rank sum test; Figure S6B). As in hM3 mice, we could reliably predict the shift in place field location for many of these simulated place cells (Figure 4B). The median prediction offset for simulated place cells was 23.0 cm (simulation n = 875, 95% CI, 20.2 – 26.0 cm), which was significantly lower than for a shuffled dataset generated by pairing the BL rate map of each cell with the CNO rate map of a randomly selected cell (see **Methods**; shuffle n = 1,204, median = 33.0 cm, 95% CI, 31.6 – 34.0 cm; simulation vs. shuffle, Z = 7.9, p = 1.3 × 10^-15^, one-sided Wilcoxon rank sum test; Figures 4C-E). After normalizing for environment size (in vivo = 60 cm diameter cylinder, simulation = 100 x 100 cm square environment), the prediction offset did not differ between simulated and experimental data (simulation, adjusted median = 12.3 cm; hM3, median = 12.1 cm; hM3 vs. simulation, Z = 0.71, p = 0.48, two-sided Wilcoxon rank sum test), demonstrating that changing the firing rates of grid subfields is sufficient to produce a predictable reorganization of hippocampal place fields.

Importantly, our simulation also pointed to a potential mechanism underlying these predictable changes in place field location. Altering grid subfield rates without adjusting grid phase preserved the spatial stability of summed grid inputs while redistributing the relative strength of its peaks, causing downstream place fields to shift to alternative, pre-existing peaks rather than random locations (Figures S6C-D). Because the simulation isolates grid subfield rates while holding all other factors constant (e.g., fixed peak firing rates, no reciprocal feedback, and no contributions from other spatial and/or directionally tuned cells), these simulations provide a direct demonstration that rate-based redistribution alone is sufficient to drive the observed place field shifts. Taken together, these results suggest that predictable hippocampal remapping can emerge from the redistribution of firing rates among spatially stable grid inputs.

### Similar grid field rate changes produce similar hippocampal remapping

To provide further support for this idea, we asked whether similar changes in grid field rates would produce a consistent reorganization of hippocampal place fields. We ran the simulation once as described above to generate a set of grid and place cells corresponding to BL and CNO1 (Figures 5A-B). Before generating a set of place cell rate maps representing CNO2, we adjusted each grid subfield rate by an amount drawn randomly from a distribution reflecting the difference in subfield rate changes between the BL and CNO sessions on each day of recording in hM3 mice (ΔBL1×CNO1 – ΔBL2×CNO2; Figures 5A-B). Since grid subfield rate changes were similar between BL and CNO sessions on consecutive days, this distribution was centered around zero (median = −0.06, 95% CI, −0.18 – −0.02).

**Figure 5.**
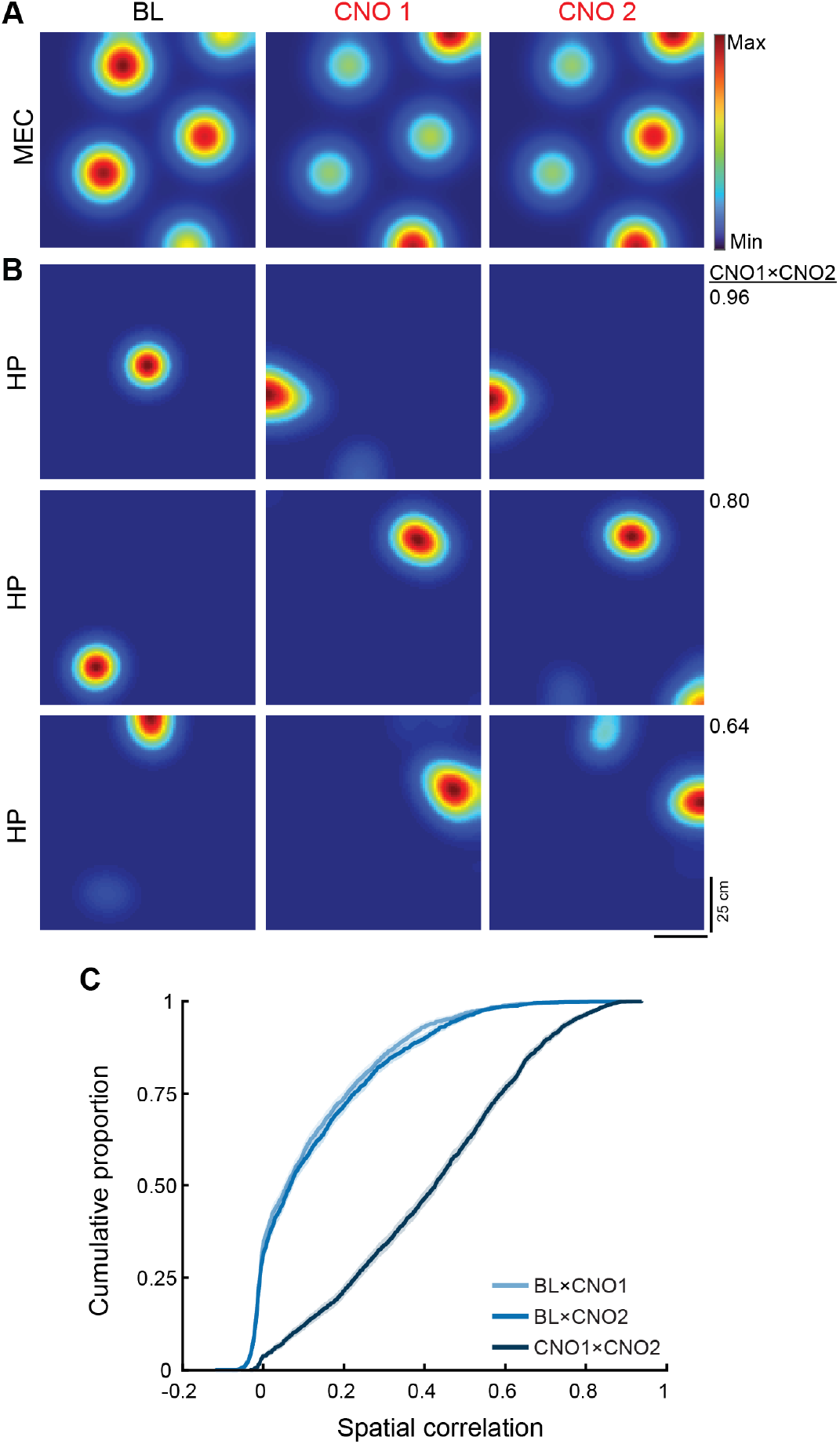
Grid subfield rate changes are sufficient to drive a reproducible reorganization of downstream place fields. **A)** Firing rate maps of one simulated grid cell before and after modification of grid subfield rates. Color indicates firing rate. We first generated a library of 10,000 grid cells with variable subfield rates (left, BL). We then modified the rate of each subfield by an amount drawn randomly from the distribution of subfield rate changes in hM3 mice (middle, CNO1). Subfield rates were then adjusted a second time by an amount drawn randomly from the distribution of rate changes observed across CNO sessions in hM3 mice (right, CNO2). **B)** Firing rate maps of three simulated place cells before and after modification of grid subfield rates (left, BL; middle, CNO1; right, CNO2). Color indicates firing rate. The spatial correlation between rate maps from CNO sessions is shown on the right for each cell. Note that all three place cells remapped in response to grid subfield rate changes (BL×CNO1 and BL×CNO2) and that the remapping was consistent between CNO1 and CNO2 (CNO1×CNO2). **C)** Cumulative distribution function shows the spatial correlation between rate maps of simulated place cells between sessions (BL×CNO1, light blue; BL×CNO2, medium blue; CNO1×CNO2, dark blue). Similar modifications to grid subfield rates produced artificial remapping on each day (spatial correlation, BL×CNO1 vs. BL×CNO2: BL×CNO1 n = 1,975, BL×CNO2 n = 1,988, D* = 0.03, p = 0.18, two-sided Kolmogorov-Smirnov test), but the reorganization of the place cell code was similar across days (spatial correlation, BL×CNO1 vs. CNO1×CNO2: BL×CNO1 n = 1,975, CNO1×CNO2 n = 2,150, D* = 0.53, p = 7.5 × 10^-253^; BL×CNO2 vs. CNO1×CNO2: BL×CNO2 n = 1988, CNO1×CNO2 n = 2150, D* = 0.51, p = 4.0 × 10^-233^; two-sided Kolmogorov-Smirnov tests).

In both runs of the simulation, there was a substantial reorganization of the place code (spatial correlation: BL×CNO1 n = 1,975, median = 0.13, 95% CI, 0.11 – 0.15; BL×CNO2 n = 1,988, median = 0.16, 95% CI, 0.14 – 0.18; Figure 5C). The degree of remapping was similar across runs (BL×CNO1 vs. BL×CNO2, D* = 0.03, p = 0.18; Figure 5C), and we could successfully use activity in the baseline session to predict place field locations during both CNO sessions (prediction offset: BL×CNO2 n = 895, median = 22.2, 95% CI, 19.4 – 24.1 cm; shuffle n = 1,218, median = 31.7, 95% CI, 30.2 – 33.1 cm; BL×CNO2 vs. shuffle, Z = 8.7, p = 4.4 × 10^-18^, two-sided Wilcoxon rank sum test; Figures 4C and S7A-C). Between CNO sessions, however, place field locations were remarkably stable, mirroring our observations in hM3 mice (spatial correlation: CNO1×CNO2, simulation n = 2,150, median = 0.67, 95% CI, 0.65 – 0.69; simulation vs. hM3, D* = 0.13, p = 0.09; Figure 5C). Together, these results demonstrate that similar changes in grid subfield rates are sufficient to drive a predictable and reproducible reorganization of downstream place fields.

## Discussion

To investigate whether the transformation from entorhinal input to hippocampal place cell output follows stable, predictable rules, we chemogenetically depolarized the same subset of MEC LII neurons across consecutive days using transgenic expression of an hM3Dq DREADD receptor. Electrophysiological recordings from MEC and CA1 revealed that artificial remapping arises from a reproducible and deterministic reconfiguration of entorhinal inputs. Repeated activation of the same subset of MEC LII neurons produced highly similar changes in grid subfield rates across days (without altering their spatial phase) and consistent changes in place cell firing patterns in CA1. This reproducibility implies that artificial remapping is constrained by preconfigured circuit dynamics, rather than stochastic processes. Moreover, we found that artificial remapping was not random: the location of a cell’s new place field could be predicted solely from its baseline activity pattern, revealing a deterministic mapping between entorhinal input and hippocampal output. Finally, by incorporating our empirically observed grid subfield rate changes into a model of the grid-to-place cell transformation, we demonstrate that consistent changes in grid field rates alone were sufficient to drive a reproducible and predictable reorganization of hippocampal place fields. Together, these results reveal that the entorhinal-hippocampal circuit can operate according to a stable input-output transformation, in which similar changes in grid subfield rates are linked to a consistent reorganization of downstream place cell activity.

Understanding the mechanisms that give rise to hippocampal remapping is crucial for explaining how the brain maintains representations of different contexts. When an animal is moved between distinct environments, place cells undergo global remapping, which is characterized by the complete orthogonalization of hippocampal activity patterns (Leutgeb et al., 2005; Alme et al., 2014; Lykken et al., 2025). This process is generally thought to arise from the coincident, independent realignment of grid modules, which decorrelates the spatial input to the hippocampus (Fyhn et al., 2007; Monaco and Abbott, 2011; Lykken et al., 2025). Under other experimental conditions, however, place cells exhibit partial remapping, meaning that some cells change firing rate and/or location while others remain stable (Shapiro, 1997; Tanila, 1997; Knierim, 2002; Anderson and Jeffery, 2003). Crucially, varying degrees of remapping have been observed without any change in grid phase (Kanter et al., 2017; Diehl et al., 2017; Lykken et al., 2025), suggesting that another mechanism is playing an important role. Our data suggest that one such mechanism may involve changes in grid subfield firing rates – a process that could reconfigure the spatial input to the hippocampus while maintaining a fixed coordinate system. By incorporating information in the firing rates of individual grid subfields, the network could maintain a stable metric for navigation while flexibly encoding multiple, context- or task-dependent representations – even within the same physical environment – substantially increasing its overall representational capacity.

In line with this, several recent studies have shown that variability among grid subfield rates exceeds chance levels and that subfield rates remain stable across sessions separated by tens of minutes (Kanter et al., 2017; Diehl et al., 2017; Ismakov et al., 2017; Dunn et al., 2017; Redman et al., 2025). Here, we extend these observations to sessions separated by more than twelve hours. In our experiments, the magnitude of grid subfield rate changes between baseline sessions on consecutive days did not differ between hM3 and control mice, indicating that within a given context, grid subfield rates remain stable over the same timescales as downstream hippocampal representations. In contrast, minor contextual changes to the environment (i.e., changes in arena shape, color, size, odor, or local/distal cues) can induce a redistribution of grid subfield rates (Diehl et al., 2017; Ismakov et al., 2017; Butler et al., 2019; Lykken et al., 2025), which has been linked to changes in both place cell firing rates (i.e., rate remapping) (Diehl et al., 2017) and place field locations (i.e., partial remapping) (Lykken et al., 2025). By comparison, our chemogenetic manipulation of MEC LII activity produced substantially larger changes in grid subfield rates than those observed in response to natural contextual manipulations, along with robust changes in both the firing rate and spatial location of CA1 place fields (Kanter et al., 2017). Therefore, a key difference between contextual versus chemogenetic manipulations may be in the extent of grid subfield rate changes, raising the possibility that the magnitude of subfield rate change determines the degree of hippocampal remapping, particularly under conditions that do not alter the spatial phase of grid cells, as is the case in our model (Figure S6B). Grid subfield rate changes may even play an important role in driving the complete orthogonalization of place cells observed between distinct environments, although this remains challenging to test directly due to coincident changes in grid phase that prevent tracking of individual subfields across environments. Additionally, it is important to acknowledge that the recurrent connectivity within the entorhinal–hippocampal circuit prevents us from providing direct, causal evidence that grid subfield rate changes elicit place cell remapping in vivo. In fact, prior computational models have suggested that variability among grid field rates arises instead from reciprocal connectivity from place to grid cells (Dunn et al., 2017; Agmon and Burak, 2020). However, the results of our feedforward grid-to-place cell simulation, which does not incorporate connections from place to grid cells, support the interpretation that changes in grid subfield rates are indeed sufficient to drive the observed reorganization of place cell activity.

If grid subfield rate changes serve as an alternative mechanism that can drive hippocampal remapping, any changes to the pattern of subfield rates – whether elicited by natural contextual changes or experimental perturbations – must be sufficiently stable to reliably support retrieval of the corresponding hippocampal representation. Our experimental and computational results support this view: similar changes in grid subfield rates across days were associated with a consistent reorganization of downstream place cell activity. This reproducibility suggests that the hippocampus does not simply respond to spontaneous fluctuations in grid cell activity, but instead transforms structured, repeatable patterns of grid cell input into distinct spatial outputs. More broadly, these findings imply that the entorhinal-hippocampal circuit may use grid subfield rate changes as a flexible but reliable signal for selecting among partially overlapping hippocampal maps. Under this view, small shifts in grid subfield rates – such as those elicited by minor contextual changes – would produce modest, partial remapping, whereas large-scale perturbations, like those induced via chemogenetic activation of MEC LII (or potentially the introduction to a novel environment), would yield a more extensive reorganization of place fields.

In many of these cases, grid subfield rate changes could act by reshaping an existing place cell representation, rather than creating a new one. Consistent with this, our empirical and computational results demonstrate that grid subfield rate changes are associated with predictable changes in place field location. In line with a previous optogenetic perturbation of CA1 neurons (McKenzie et al., 2021), we found that CA1 place cells frequently shifted the location of their fields toward positions with sparse, but detectable activity instead of random locations, indicating that the underlying map was reorganized rather than replaced. In our grid-to-place cell simulation, we observed that the spatial stability of total grid input was also maintained following changes in subfield rates, indicating a similar redistribution of activity among underlying grid inputs. In sum, we demonstrated that place cell remapping can emerge from the coordinated redistribution of grid input that results from population-level changes in grid subfield rates. Under these conditions, place cell remapping may be constrained by intrinsic network dynamics that guide the reorganization of spatial representations toward locations with preexisting spiking activity, rather than random locations.

Taken together, our results demonstrate that CA1 place cell activity reorganizes in a reproducible and predictable manner following the chemogenetic depolarization of a subset of MEC LII neurons on consecutive days, indicating that the entorhinal-hippocampal circuit implements a stable and deterministic input-output transformation. In line with our modeling results, consistent alterations in grid subfield firing rates observed across days provide a plausible mechanism for reproducible and predictable remapping in CA1, even in the absence of changes in grid phase. The reproducible reorganization of place fields across days implies that artificially induced remapping is constrained by the underlying anatomical connectivity of the circuit, despite the dense, recurrent architecture of CA3. The predictable nature of the hippocampal response suggests that, under these conditions, place cell remapping reflects preconfigured network dynamics operating on an existing spatial map. This form of remapping is therefore distinct from global remapping, which involves the orthogonalization of hippocampal activity patterns and the apparently random reassignment of place field locations. More broadly, our results support an alternative mechanism by which the hippocampus can update spatial representations without requiring independent changes in the spatial phase of grid modules. Within this framework, the network may flexibly employ these rate-based and/or phase-based mechanisms depending on the magnitude of contextual change. Minor contextual changes are likely to produce modest adjustments in grid subfield rates and partial remapping of place cells, whereas major contextual changes are more likely to engage phase-based mechanisms, with or without accompanying changes in grid subfield rates. Future work will be essential for determining how these mechanisms interact during natural behavior and how they contribute to context discrimination, memory retrieval, and navigation in complex environments.

## ACKNOWLEDGEMENTS

We thank Qiangwei Zhang for histology and Edvard Moser for comments on the manuscript. Supported by NIH Grant RO1MH097130-05, Kavli Foundation, the Centre of Excellence scheme of the Research Council of Norway – Centre for Neural Computation (Grant 223262/F50) and Center for Algorithms in the Cortex (Grant 332640), Research Council of Norway “Toppforsk” Grant 249945, Marie Curie Grant MGATE (Grant 765549), the Trond Mohn Foundation (Grant TMS2021TMT04) and the National Infrastructure scheme of the Research Council of Norway – NORBRAIN (Grant 295721).

## AUTHOR CONTRIBUTIONS

C.M.L., B.R.K., J.K. and C.G.K. planned and designed experiments, conceptualized and planned analyses, and interpreted data; C.M.L., B.R.K., J.K., O.M.T.C., K.A., and L.A.L.D. performed experiments; C.M.L., B.R.K. and L.A.L.D. visualized and analyzed data; C.M.L. curated the data; C.M.L. wrote the first draft of the paper with periodic input from B.R.K.; C.M.L. and L.A.L.D. edited the paper with periodic input from B.R.K. and C.G.K. All authors discussed the data. C.G.K. supervised and funded the project.

## COMPETING FINANCIAL INTERESTS

The authors declare no competing interests.

## Methods

### Experimental Model and Subject Details

Our subjects were adult (2 - 6 months, 17 – 37 g) male and female mice. We crossed the EC-tTA line (Mutant Mouse Resource & Research Centers, Stock: 031779-MU; RRID: MMRRC_031779-MU) to an hM3Dq-tetO line (Jackson Laboratory, Stock: 014093; RRID: IMSR_JAX:014093) to enable control of neurons in the superficial layers of medial entorhinal cortex. Pups were evaluated for transgene expression via PCR of genomic DNA isolated from tail biopsies. For hippocampal and MEC recordings, the hM3 group consisted of Ent/tTA x hM3-DDD double-positive offspring. The hippocampal and MEC control groups included hM3 mice injected with saline and littermate controls injected with either CNO or saline. The hippocampal control group includes a single C57BL/6J (Jackson Laboratory, Stock: 000644; RRID: IMSR_JAX:000664) mouse injected with saline. The MEC control group also includes a single C57BL/6J (Jackson Laboratory, Stock: 000644; RRID: IMSR_JAX:000664) mouse injected with saline. Mice were kept on a 12-hr light/dark schedule and were fed ad libitum. They were housed in environmentally-enriched transparent Plexiglas cages in a humidity- and temperature-controlled environment. Mice were group-housed prior to surgery and then housed separately. All procedures were approved by the Institutional Animal Care and Use Committee at the University of Oregon and the National Animal Research Authorities of Norway. They were performed according to the Norwegian Animal Welfare Act and the European Convention for the Protection of Vertebrate Animals used for Experimental and Other Scientific Purposes.

### Histological Procedures

When electrophysiological recordings were complete, mice were administered a lethal dose of pentobarbital sodium (Euthasol, 50 mg/kg) and perfused transcardially with 4% paraformaldehyde in phosphate-buffered saline (PBS). After an additional 24 hours of post-fixation in paraformaldehyde overnight, the brains were extracted and transferred to a 30% sucrose solution. For identification of recording sites, brains were sectioned at 30 μm, mounted on glass microscope slides, and the tissue was stained with Cresyl violet. The slides were then coverslipped and examined under the microscope.

For identification of the hM3Dq transgene localization, free-floating 30 μm thick sections were collected for immunohistochemistry. As hM3Dq has a HA tag, we performed an anti-HA stain to identify the transgene and an anti-NeuN counterstain to delineate layers in MEC. Briefly, sections were washed in PBS, permeabilized using PBS containing 1% of Triton (Merck), and blocked in PBS containing 10% of normal goat serum (Abcam) and 1% of Triton. Primary antibody incubation using a rat anti-HA antibody (1:500 dilution factor, reference11867423001, Roche) and guinea pig anti-NeuN antibody (dilution factor 1:1000, reference ABN90P, Millipore) was done for 48 hours at 4°C. Sections were then washed in PBS and underwent secondary antibody incubation using goat anti-rat Alexa Fluor 546 (dilution factor 1:400, reference A11081, Invitrogen) and goat anti-guinea pig Alexa Fluor 647 (dilution factor 1:400, reference A21450, Life Technologies) for 2 hours at room temperature. Sections were then mounted on glass microscope slides, then once dried cleared for 10 min in Toluene (BDH Prolabo) and coverslipped using a mixture of Toluene and Entellan (Merck). Fluorescence images were obtained using a slide scanner (Axio Scan.ZI, Zeiss) and a confocal microscope (LSM 880, Zeiss).

### Surgical Procedures

All surgeries were performed using aseptic techniques on experimentally-naive mice. Prior to surgical implantation of the microdrives, ketamine (100 mg/kg) was administered as a preanesthetic. Dexamethasone (0.1 mg/kg) and atropine (0.03 mg/kg) were also administered presurgically to ameliorate possible inflammation and respiratory irregularities, respectively. Surgical anesthesia was maintained with isoflurane (1.25%–2.0%, adjusted as necessary for appropriate depth of anesthesia). Eyes were moistened with antibacterial ophthalmic ointment. Mice were placed in a stereotaxic frame and held in position with atraumatic ear bars. The skull was exposed and lambda and bregma were zeroed in the vertical plane. The surface of the skull was cleaned with hydrogen peroxide, lightly scored with a scalpel blade, and coated with a thin layer of cyanoacrylate glue that was allowed to dry completely before proceeding. For recordings from CA1, one craniotomy was drilled in the left hemisphere overlying the dorsal hippocampus (centered at AP: −1.8 mm; ML: 1.2 mm relative to bregma). For recordings from medial entorhinal cortex (MEC), one craniotomy was drilled in the left hemisphere, exposing the transverse sinus 3.4 mm lateral to the midline. Four to six additional holes were drilled around the perimeter of the skull for stainless steel anchor screws (00–90 x 1/8”) and ground wires from the recording array. The tetrodes of the array were lowered into the cortex overlying the hippocampus or MEC to a depth of approximately 0.65 mm. In MEC, the tetrode array was implanted 300-500 mm anterior of the transverse sinus at a 3-6 degree angle aimed posteriorly. After the tetrodes were in place, sterile Vaseline was applied to isolate the tetrodes, preserving the ability to adjust tetrode depth. Dental cement was applied to secure the array to the skull. Mice were subcutaneously administered buprenorphine (0.06 mg/kg) postoperatively for analgesia to minimize discomfort.

### Electrophysiology Protocol

All implanted mice were allowed to recover from surgery for at least seven days, after which screening for units began. A tethered HS-16 or HS-18MM operational amplifier (Neuralynx, Bozeman, MT) was plugged into the tetrode recording array to monitor/record behavior and neuronal activity. Recording sessions occurred based on the presence of neural activity, regardless of the light/dark cycle. MEC screening and recording sessions were performed in a 60 x 60 cm square environment, a 100 x 100 cm square environment, or a 90 x 120 cm rectangular environment with dominant visual cues. Hippocampal screening and recording sessions were performed in a 60 cm diameter cylinder or a 60 x 60 cm square environment with dominant visual cues. During initial screening sessions, the array was moved down 45-90 μm per day, and an audio channel was monitored for evidence of theta rhythmicity and/or the occurrence of sharp waves. Recordings of MEC activity were initiated when cells with clear spatial or head direction correlates were first observed along with increased power in the theta range. Recordings of hippocampal activity were initiated when spiking activity with clear spatial correlates was first observed along with increased power in the theta range.

For all experiments, baseline (BL) activity was recorded for 30 min. Mice were then removed from the cylinder and given an intraperitoneal injection of either clozapine N-oxide (CNO, Sigma-Aldrich, St. Louis, Missouri, USA) (hM3: 1 mg/kg, 0.1 mg/ml in 10% DMSO/saline solution) or saline. Immediately following the injection, mice were placed back into the environment. On Day 1 of experiments repeated across days (Figures 1 and 2), data were recorded during the initial BL session (BL1) and a two-hour recording session following CNO or saline injection (CNO1). On Day 2, after a 12+-hour delay, data were recorded during the subsequent BL session (BL2) and a two-hour recording session following CNO or saline injection (CNO2). Experiments were repeated for each mouse as long as activity was present.

### Single Unit Recording

Tetrodes were made by spinning together four lengths of 18-micron-diameter 10% iridium/platinum wire (California Fine Wire, Grover Beach, CA) and applying heat to fuse the polyamide coating at one end. We used custom-made four-tetrode recording arrays, VersaDrive-4 microdrives (Neuralynx), or MDR-16 four-tetrode microdrives (Axona Ltd, St. Albans, U.K.) with custom-ordered with Mill-Max (Mill-Max Mfg. Corp., Oyster Bay, NY) connectors. The custom-made four-tetrode recording arrays were adapted from previously published work (Gray et al., 1995). The coating on the free ends of each wire was removed and each uncoated wire segment was inserted into a channel of an EIB-16 electrode interface board (Neuralynx) and fixed in place with a gold-coated pin. Each EIB-16 loaded with four tetrodes was fixed to a Teflon stage mounted on three drive screws. The drive screws (0–80 x 3/8”) allowed depth adjustments of the entire array and served as a structural link to the skull. The tetrodes could each be lowered independently when VersaDrive-4 microdrives were used. Neuronal data were acquired using the Cheetah-16 system and Digital Lynx 4SX systems (Neuralynx). Recorded signals were amplified automatically for each tetrode when the experimenter selected an appropriate input range (typically ± 250-800 mV). The signals were band-pass filtered (spikes: 600 – 6000 Hz; local field potential: 0.1 – 475 Hz) and stored using Neuralynx data-acquisition software.

Thresholds were set such that only waveforms of a specified minimum voltage (e.g., 50 μV) were stored. A digital camera mounted above the recording environment and linked to the Cheetah-16 system recorded the position of the mouse by tracking two light-emitting diodes fixed to the headstage and aligned with the body axis of the mouse.

### Unit Isolation and Recording Stability

Unit isolation and assessment of recording stability of MEC and CA1 recordings were performed on 30- or 60-min epochs. For both Day 1 and Day 2 of repeated experiments, we used the 30-min BL sessions (BL1, BL2) and the two-hour post-injection sessions (CNO1, CNO2), which were either divided into 30-minute or 60-minute epochs for analysis purposes. Units were manually separated offline with MClust spike-sorting software (courtesy of David Redish, University of Minnesota) for MATLAB (MathWorks, Natick, MA) using the previously described standards for unit isolation (Kentros et al., 2004). Cluster boundaries were applied across successive sessions to track clusters over time. Isolated clusters corresponding to putative pyramidal neurons formed clear Gaussian ellipses generally based upon peak-to-peak projections of different tetrode wires with minimal overlap with neighboring clusters or noise. These clusters were divided into one of three groups according to a subjective judgment of quality (Q), as described previously (Kanter et al., 2017). Q-1 clusters had virtually no overlap on at least one projection and no events within a 2 ms refractory period; Q-2 clusters included clear Gaussians with a small degree of overlap with other clusters or noise; Q-3 cells met neither criteria; Q-off cells did not have enough spikes to judge the quality. Neurons categorized as Q-3 were not included for any analyses. Putative interneurons with generally spherical clusters were assigned Q-values exclusively by cluster boundary criteria. Cluster boundaries were then applied across successive epochs and minor adjustments were made when necessary to optimally separate clusters from each other and from noise. Inspection of spike waveforms, inter-spike intervals, autocorrelations, and cross-correlations were used as additional methods to ensure each cluster was correctly tracked over time. In two-session comparisons (i.e., spatial correlations, difference scores, etc.), it was required that clusters in both sessions passed criteria to be included. In CA1, we recorded 778 cells from 36 mice (19 hM3+ mice injected with CNO, 12 hM3-mice injected with CNO, 9 hM3+ mice injected with saline, and 1 C57BL/6J injected with saline) that met our standards of cluster quality. In MEC, we recorded 310 cells from 13 mice (4 hM3+ injected with CNO or saline, 4 hM3+ mice injected with CNO, 4 hM3-mice injected with CNO, and 1 C57BL/6J injected with saline) that met our standards of cluster quality.

### General Electrophysiological Analysis

In order to exclude spiking activity occurring during periods of immobility, a walk filter (*≥* 2 cm/s) was applied. Rate maps were then generated by binning the location of each spike for each epoch, dividing the number of spikes in each bin by the time spent in that bin, and smoothing with a Gaussian. For 60 cm diameter cylindrical and 60 x 60 cm square environments, 2 x 2 cm bins were used. For 100 x 100 cm square and 90 x 120 cm rectangular environments, 4 x 4 cm bins were used. Mean firing rate was defined as the total number of spikes divided by the duration of the recording session. Peak firing rate was defined as the maximal firing rate of all spatial bins. To assess spatial correlation, pairs of rate maps were each reshaped into a single vector and the correlation coefficient (Pearson’s linear correlation) between these vectors was calculated. Pixels of incongruity between the two vectors, resulting from unvisited pixels in either epoch, were excluded from the calculation. To generate a shuffled control group of spatial correlation values, we calculated the correlation coefficient between the rate map of each cell in the first session and the rate map of a randomly selected cell in the second session. Difference scores (i.e., normalized change) were calculated as: (session 2 value – session 1 value) / (session 2 value + session 1 value). These scores are reported in Figures 1, S3 and S4. We reported the absolute value of these difference scores in Figures S3, S4, and S6.

### Functional Classification of Place Cells

Place cells were defined as putative excitatory neurons (mean firing rate < 7 Hz) with high spatial stability (spatial correlation between first and second halves > 0.5) and/or high spatial information content in the BL session, peak rate > 0.5 Hz, and at least one identified place field. Place fields were defined as areas with at least 10 contiguous pixels (40 cm^2^) where the firing rate exceeded 40% of the peak rate. Cells were required to meet all defined criteria in either the baseline or the CNO session to be included. For repeated experiments across days, place cells were also required to be stable across baseline sessions. We used either the 30-60 or the 90-120 min epochs of the two-hour post-injection recording session on Day 1 and Day 2 to capture the peak activity of CNO or to examine the activity at the end of this two-hour period, respectively.

### Predictability Analyses

For predictability analyses, we focused on the first 30-min epoch after CNO injection in order to capture the first location that CA1 place fields shifted to after CNO injection. For each cell, we first identified the location of the primary place field in the BL and CNO sessions. We restricted our analysis to cells with an identified place field that were active in both sessions and exhibited large changes in the place field location between sessions. To define a threshold for classifying whether place fields shifted between sessions, we generated a random reference distribution with mean μ = 30 cm and standard deviation σ = 6 cm. The 95th percentile of this distribution (20 cm) served as our lower bound for determining whether place field locations shifted between sessions cells.

For each of the place cells that shifted between, we then identified up to three regional maxima from the rate map of the BL session that served as predictions for future place field locations. We calculated the distance between each of these locations and the location of the primary place field in the CNO session, defining the shortest of these distances as the ‘prediction offset’. We used the same process to predict the location of place fields in a separate group of mice moved between two distinct, familiar environments (A and B). To generate a shuffled control group to compare with our empirical data in hM3 mice (Figure S5B), we identified up to three regional maxima from the rate map of the BL session for each cell. We then calculated the distance between each of these predicted locations and the location of the primary field in the CNO session for a randomly selected cell, defining the shortest of these distances as the ‘prediction offset’.

### Functional Classification of MEC Cells

For repeated experiments across days, we used either the 30-60 or the 30-90 min epochs of the two-hour post-injection recording session on Day 1 and Day 2 to capture the peak activity of CNO and ensure that the coverage of the environment was complete. Cells with a mean firing rate < 10 Hz were classified as putative excitatory neurons. Firing fields were defined as areas of at least 80 cm^2^ where the firing rate exceeded 20% of the peak rate. Firing fields with peak rates lower than 1 Hz were ignored.

Grid cells were identified by calculating a spatial autocorrelation map for each unsmoothed rate map (Sargolini et al., 2006). A cell’s spatial periodicity was determined by comparing a central circular region of the autocorrelogram, excluding the central peak, with versions of this region rotated at 30°increments (Sargolini et al., 2006; Langston et al., 2010). Pearson correlations were calculated by comparing the circular region to all rotated versions. 60°and 120°rotations should have high correlation scores due to the triangular pattern of the grid, whereas 30°, 90°, and 150°rotations should result in low correlations. Therefore, a cell’s grid score was defined as the minimum difference between correlation scores for either rotation from the first group and any rotation in the second group (range: –2 to 2). Cells with grid scores exceeding the 95^th^ percentile of a shuffled distribution were classified as grid cells. To generate the shuffled distribution, the spikes of each cell were circularly shifted in time relative to the mouse’s position by a random amount between 20 s and 20 s less than the total length of the recording session. The grid score was then calculated using these shuffled spike times, and this procedure was repeated 500 times for that cell. A distribution of values was generated including the 500 shuffled results from all cells, and the 95th percentile of this shuffled distribution was used as the cutoff for classifying grid cells.

### Analysis of Individual Grid Subfields

Individual grid subfields were defined as areas of at least 40 cm^2^ where the firing rate exceeded 20% of the peak rate. Fields with peak rates lower than 0.5 Hz were ignored. Individual subfield rates were defined as the peak rate of each identified firing field. Changes in grid subfield rates between sessions were assessed using absolute difference scores. The correlation between grid subfield rate changes across days (i.e., Figure 1C) was obtained by calculating the Pearson correlation between vectors representing the change in peak firing rate for each subfield between the BL session to the CNO session on Day 1 and Day 2. To create a shuffled control group, for each subfield, we calculated the change in peak firing rate from the BL session to the CNO session on Day 1. Then, we calculated the change in the peak firing rate of that field in the BL session on Day 2 and the peak firing rate of a randomly selected subfield in the CNO session on Day 2. A grid field rate change correlation for this shuffled control group was obtained by fitting a line between points representing the change in peak field rates on Day 1 and Day 2 (Figure 1C).

To quantify the relationship among subfields for each grid cell in each session, we ordered subfield firing rates for each grid cell from highest to lowest. We then calculated the normalized difference in peak firing rate between all subfield pairs and arranged these values into a single vector representing grid subfield relationships in that session. To test whether grid subfield relationships were maintained across sessions, we calculated the Pearson correlation between vectors representing subfield relationships in each session (Figure S3F). The r value (coefficient of correlation) and p value are reported for the fit.

### Model of Grid-to-Place Cell Transformation

To evaluate the impact of changing the firing rates of grid subfields on hippocampal place cells, we applied empirically determined grid subfield rate changes to a competitive linear summation model of the grid-to-place cell transformation (de Almeida et al., 2012). Briefly, this model is a three-layer network where place fields are created by linear summation of weighted inputs from entorhinal grid cells and dentate gyrus granule cells. Competitive interactions limit the active pool of neurons to only those receiving the most excitation (10% winner-take-all process), which is meant to mimic gamma frequency feedback inhibition.

To generate a library of 10,000 simulated grid cells with variable rates across their subfields (e.g., Figure 4A, left), we modified the subfield rates of grid cells from the original model (grid scale logarithmically distributed between 30 and 100 cm, one of three random orientations, random spatial phase). Grid subfield centers were identified as described above and used as seed locations for new grid cell maps. Each seed location received a peak firing rate randomly sampled from a normal distribution (μ = 12 Hz, σ = 3 Hz). Rate maps were then smoothed with a Gaussian kernel to create grid subfields, where the kernel size ranged from 5 to 12.5 spatial bins depending on the number of grid fields in the map. Finally, rate maps were normalized between 0 and 1 Hz (matching the original model) to isolate the effect of changing subfield rates while and exclude the effect of variation in the overall peak firing rate of grid cells. These grid cells with variable subfield rates represented the BL session and a set of simulated place cells were created from these grid cells as in the original model.

From our empirical data, we created a distribution of rate difference scores by calculating the change in peak firing rate for each grid subfield between BL and CNO sessions (Figure S6A). We applied these empirically determined subfield rate changes to our simulated grid cells by adjusting the firing rate of each subfield by an amount randomly selected from this distribution (e.g., Figure 4A, right). These grid cells represented the CNO session and a second set of rate maps for the simulated place cells were created from these modified grid cells as in the original model.

The simulation was run with the following parameters: arena size = 100 x 100 cm, bin size = 1 cm, number of grid cells = 10,000, number of granule cells = 200, E = 10%, relative DG-to-MEC input to CA3 (R) = 0.24, place field rate threshold = 20%, minimum place field size = 200 cm^2^. Note that all connections and weights were held constant from the BL simulation to isolate the effect of grid subfield rate changes.

For each simulated place cell, we evaluated whether we could predict the location of its place fields in the CNO session using the rate map from the BL session using the same method described for our empirical data from hM3 mice. To generate a shuffled control dataset for comparison, we identified up to three regional maxima from the rate map of the BL session for each cell. We then calculated the distance between each of these predicted locations and the location of the primary place field in the CNO session for a randomly selected cell, defining the shortest of these distances as the prediction offset.

To evaluate the extent of reorganization among grid inputs following subfield rate changes, we generated excitation maps representing the summed grid cell input to each simulated place cell before and after grid field rates were modified (e.g., Figure S6C). Although the model incorporates both direct and indirect grid input to the hippocampus, we limited our analysis to the direct grid inputs under the assumption that subfield rate changes would similarly affect all input streams. To quantify the reorganization among grid inputs between sessions, we first calculated the spatial correlation between excitation maps representing the BL and CNO sessions (Figure S6D). Then, using the same method as for place cells (Figure 3), we assessed whether the location of peaks within each excitation map shifted to a predictable location (i.e., an existing peak in the excitation map in the BL session) after grid field rates were modified. Fields in the summed excitation maps were defined as areas where the firing rate exceeded 50% of the peak rate, which was required to be > 0.3 Hz. For the shuffled control dataset, the excitation map representing summed grid inputs to each simulated place cell in the BL session was used to generate up to three predictions of future peaks among summed grid inputs. We then calculated the distance between each of these predicated locations and the observed location of peak firing in the excitation map of a randomly selected simulated place cell in the CNO session.

After the first run of the simulation (BL and CNO1), we adjusted the firing rates of grid subfields a second time, by amounts we observed empirically in hM3 mice. To do this, we calculated the normalized difference in peak firing rate between the BL and CNO session for each grid subfield in hM3 mice on both recording days (as in Figure 1C; Day 1: Δ BL1×CNO1; Day 2: Δ BL2×CNO2). We then generated a distribution reflecting the difference between subfield rate changes on each day by subtracting the distribution for Day 2 from the distribution for Day 1 (Δ BL1×CNO1 – Δ BL2×CNO2). The fact that this distribution was centered around zero (median = −0.06, 95% CI, −0.18 – −0.02) reflects that subfield rate changes were similar between BL and CNO sessions across days. We then adjusted the firing rate of each simulated grid subfield by an amount that was randomly selected from this distribution before generating a second set of simulated place cells, corresponding to CNO2 (Figures 5A-B). Again, all variables in the simulation were held constant except grid subfield rates.

Finally, a separate simulation was run to evaluate the degree of reorganization among grid inputs following the independent realignment of grid cells in different modules (Figure S6C). Grid cells had subfields with uniform peak firing rates. Rather logarithmically distributing grid spacing between 30 and 100 cm (as in the original model), we created three grid modules consisting of simulated grid cells with spacings of 30, 50, or 100 cm (30 cm, n = 5,500; 50 cm, n = 3,000; 100 cm, n = 1,500). Grid cells within each module had the same orientation, and module orientations were offset by 60°. These grid cells represented the BL session and a set of simulated place cells were created from these grid cells as in the original model. Each grid module was then shifted randomly (x and y offsets sampled independently) by an amount between 0 cm and half of the grid spacing for that module. Note that all grid cells within a module shifted coherently, but there was no relationship between the phase changes of different modules. These grid cells represented the CNO session and place cells were created from these grid cells as in the original model. We quantified the extent of reorganization among grid inputs in this simulation as described above for our simulation grid subfield rate changes.

### Quantification and Statistical Analysis

Unless otherwise stated, all analyses were conducted using MATLAB (MathWorks). The experimenter was blind to the mouse’s genotype and experimental grouping during unit isolation. Two-sided statistical tests were used for post hoc analyses and one-sided tests were used when there was a clear a priori prediction. Statistical significance was defined with alpha level = 0.05. Nonparametric tests were used when the assumptions for parametric tests were clearly violated. Median values are reported/displayed for nonparametric tests and mean values are reported/displayed for parametric tests. Error is reported using the 95% confidence interval (95% CI) or standard error of the mean (SEM). Smoothed distributions of the data were estimated using kernel density estimation.

Two-group estimation plots were generated using the DABEST Python package (Ho et al., 2019). In these plots, the black dot indicates the median difference, the black bars are error bars depicting 95% confidence intervals, and the filled curve is the bootstrapped sampling-error distribution.

## Data and Software Availability

All quantification methods used in the custom scripts are described above. Further requests for custom scripts and data used in this study should be directed to the corresponding author and will be fulfilled upon reasonable request.

## Supplementary Figures

**Figure S1.**
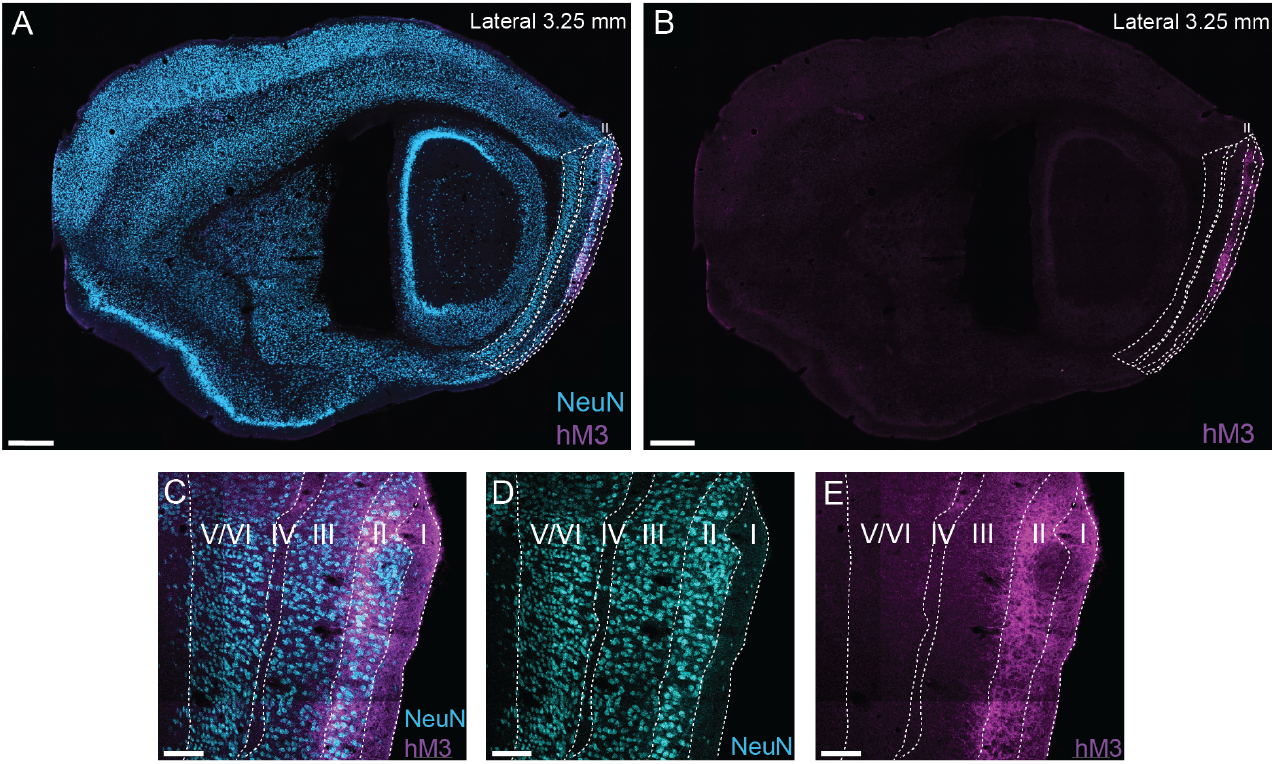
Transgenic expression of hM3Dq DREADD receptor in MEC LII. **A)** Representative image of fluorescent immunohistochemistry of a sagittal brain slice targeting the transgene hM3Dq (magenta). Expression is largely restricted to entorhinal cortex layer II. Counterstain NeuN (cyan). Scale bar: 500 μm. **B)** Same as in (A) without the NeuN counterstain. **C-E)** Representative high magnification images of dorsal MEC showing NeuN (cyan) and hM3 (magenta) staining. Cropped images from the same brain-wide section shown in (A) and (B). Scale bar: 100 μm.

**Figure S2.**
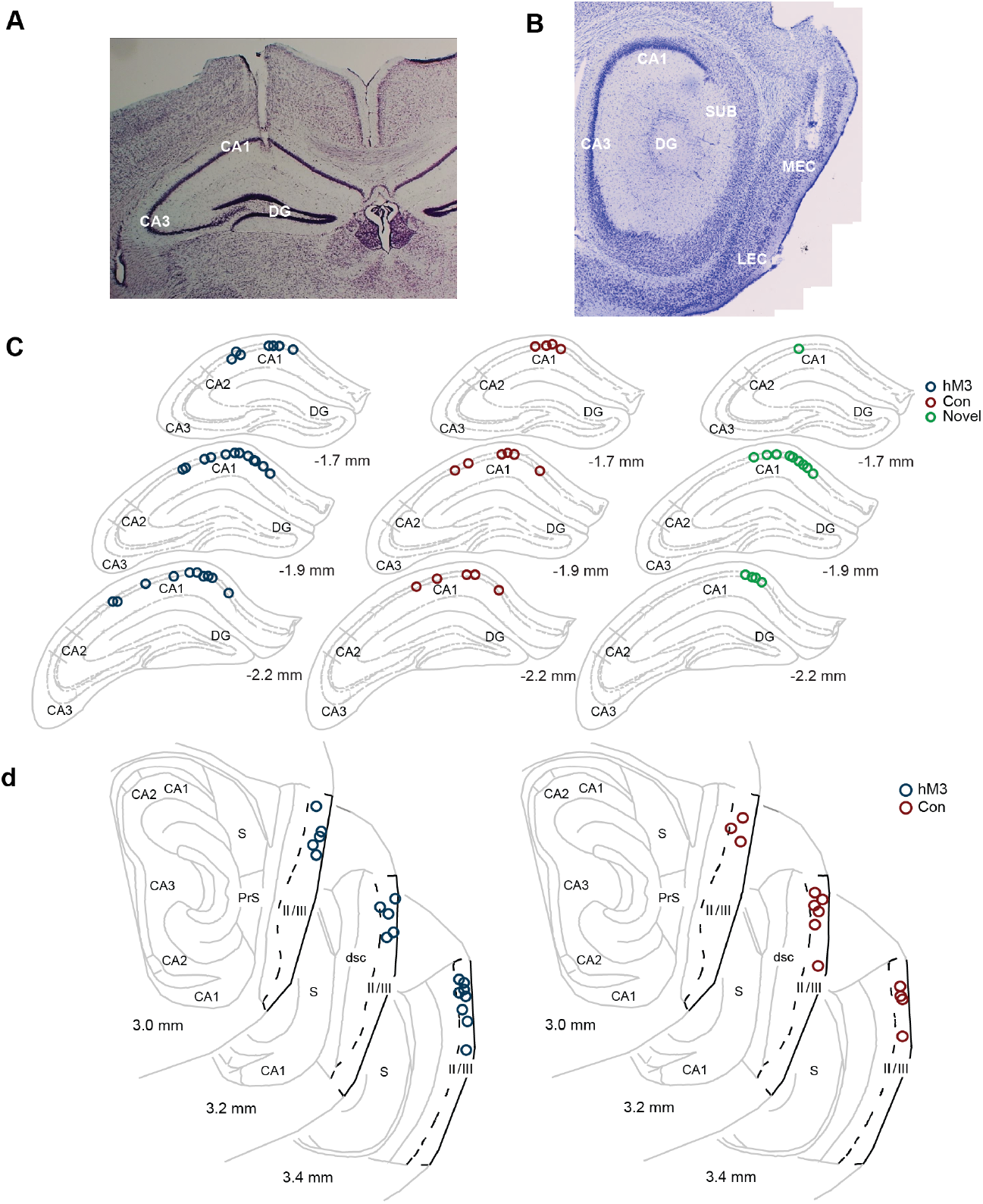
Recording sites in CA1 and MEC. **A)** Representative coronal section used to identify tetrode tracks in CA1. **B)** Representative sagittal section used to identify tetrode tracks in superficial MEC. **C)** Tetrode locations in CA1 identified in three coronal sections in hM3 mice (left, blue), control mice (middle, red), and a separate group of mice exposed to two distinct environments (right, green). Numbers indicate distance from bregma. **D)** Tetrode locations in superficial layers (II/III) of MEC identified in three sagittal sections in hM3 (left, blue) and control (right, red) mice. Numbers indicate distance from midline. SUB, subiculum; PrS, presubiculum; dsc, lamina desiccans.

**Figure S3.**
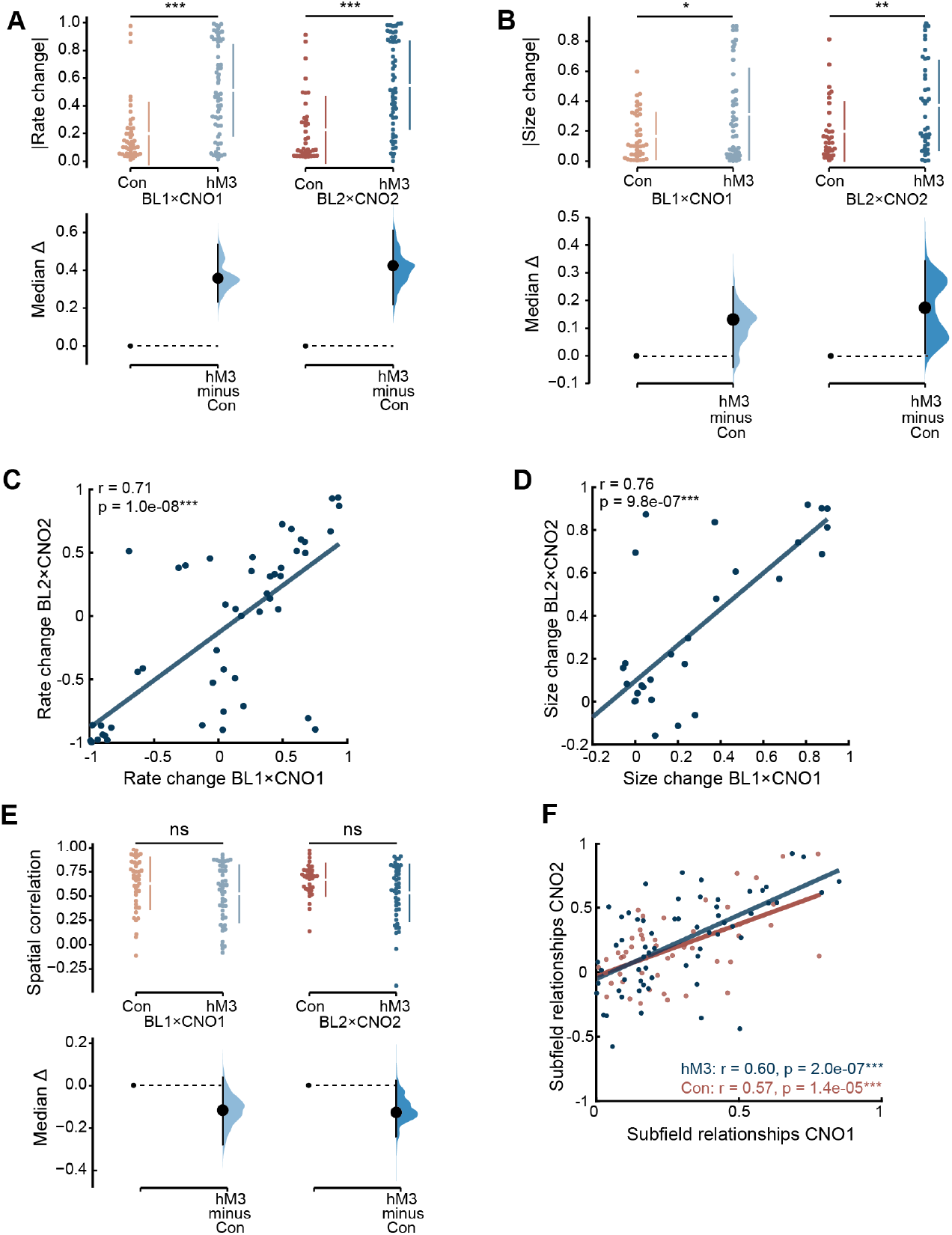
Putative excitatory neurons in MEC exhibit similar changes in firing rate and field size across days. **A-B)** Panels show significant changes in firing rate (A, left) and field size (B, right) of putative excitatory neurons in MEC between BL and CNO sessions on Day 1 and Day 2 in hM3 (blue) versus Con (red) mice (rate change, hM3 vs. Con: BL1×CNO1, hM3 n = 61, Con n = 46, Z = 4.8, p = 8.8 × 10^-7^; BL2×CNO2, hM3 n = 58, Con n = 39, Z = 5.0, p = 3.3 x10^-7^; size change, hM3 vs. Con: BL1×CNO1, hM3 n = 46, Con n = 42, Z = 1.7, p = 0.04; BL2×CNO2, hM3 n = 41, Con n = 33, Z = 2.5, p = 6.6 × 10^-3^; one-sided Wilcoxon rank sum tests). Points represent individual MEC neurons; gapped lines represent mean ± standard deviation. Change refers to an absolute difference score (see **Methods**). **C-D)** Scatterplots show significant correlation between firing rate (C, left) and field size (D, right) changes of putative excitatory neurons in MEC between BL and CNO sessions on Day 1 and Day 2 in hM3 mice (rate change, BL1×CNO1 vs. BL2×CNO2: n = 49, r = 0.71, p = 1.0 × 10^-8^; size change, BL1×CNO1 vs. BL2×CNO2: n = 30, r = 0.76, p = 9.8 × 10^-7^; linear correlations). Points represent individual MEC neurons. *\*\*\*p* < 0.001. **E)** Panel shows that there was no difference in the spatial correlation of MEC neurons (including grid cells) following CNO injection between hM3 and control mice (spatial correlation, hM3 vs. Con: BL1×CNO1, hM3 n = 48, Con n = 42, Z = 1.6; p = 0.10; BL2×CNO2, hM3 n = 43, Con n = 34, Z = 1.8; p = 0.07; two-sided Wilcoxon rank sum tests). Points represent individual MEC neurons; gapped lines represent mean ± standard deviation. **F)** Scatterplot showing significant correlation between grid subfield relationships during CNO session on Day 1 and Day 2 in hM3 (blue) and Con mice (red) (CNO1 vs. CNO2: hM3 n = 66, r = 0.60, p = 2.0 × 10^-7^; Con n = 64, r = 0.57, p = 1.4 × 10^-5^; linear correlations). For each grid cell, subfield rates were ordered from highest to lowest. We then calculated the normalized difference in peak firing rate between all subfield pairs. Points represent the normalized difference between each subfield pair in CNO1 versus CNO2. *\*\*\*p* < 0.001. For lower panels in (A), (B) and (E), black dot: median; black bars: 95% confidence interval; filled curv.e: sampling-error distribution.

**Figure S4.**
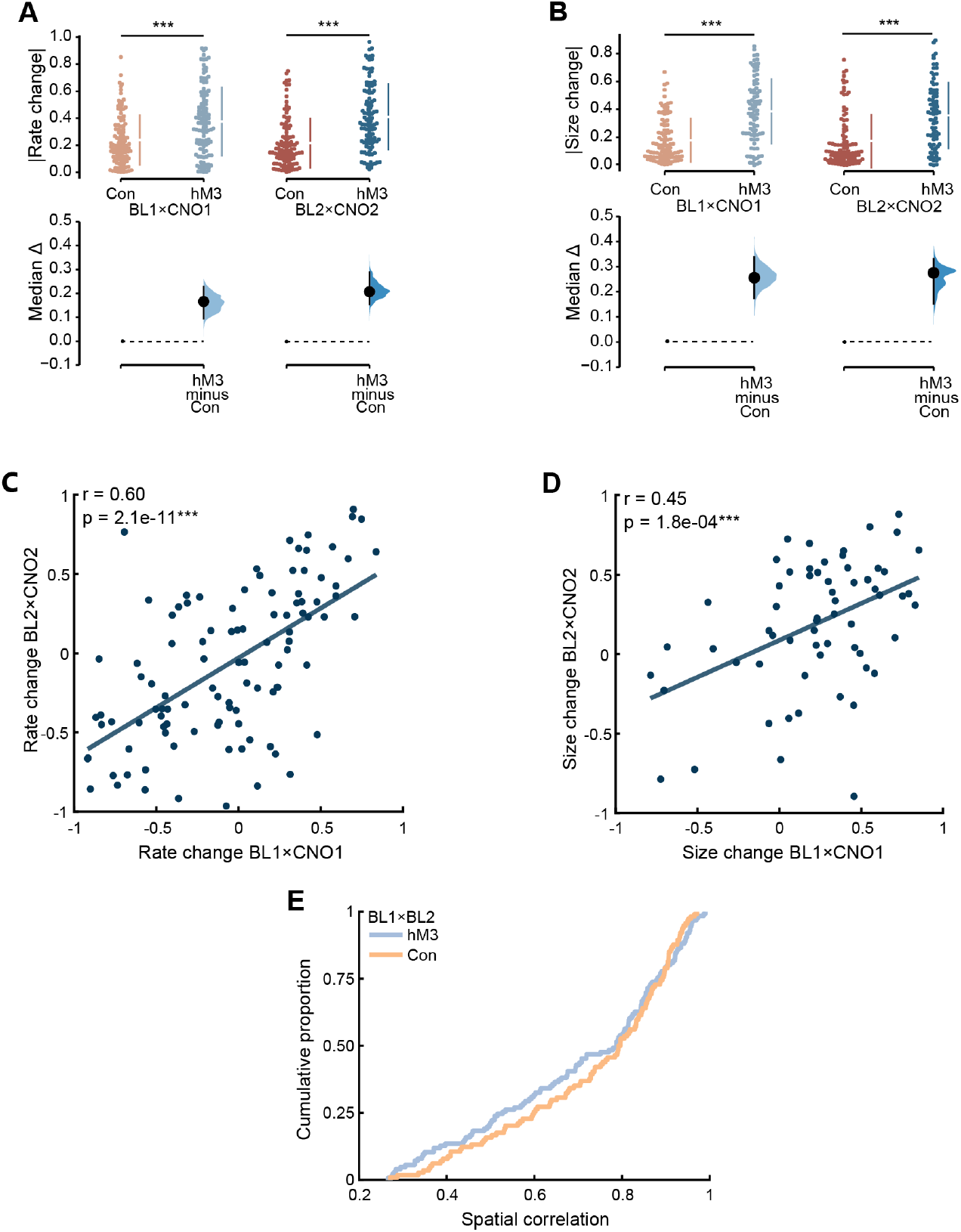
CA1 place cells exhibit consistent changes in firing rate and field size across days. **A-B)** Panels show significant changes in firing rate (A, left) and field size (B, right) of CA1 place cells between BL and CNO sessions on Day 1 and Day 2 in hM3 (blue) versus Con (red) mice (rate change, hM3 vs. Con: BL1×CNO1, hM3 n = 113, Con n = 109, Z = 4.1, p = 1.8 × 10^-5^; BL2×CNO2, hM3 n = 109, Con n = 111, Z = 6.3, p = 1.6 × 10^-10^; size change, hM3 vs. Con: BL1×CNO1, hM3 n = 86, Con n = 101, Z = 6.1; p = 4.6 × 10^-10^; BL2×CNO2, hM3 n = 80, Con n = 102, Z =5.5, p = 2.0 × 10^-8^; one-sided Wilcoxon rank sum tests). Points represent CA1 place cells; gapped lines represent mean ± standard deviation. Change refers to an absolute difference score (see **Methods**). **C-D)** Scatterplots show significant correlation between firing rate (C, left) and field size (D, right) changes of CA1 place cells between BL and CNO sessions on Day 1 and Day 2 in hM3 mice (rate change, BL1×CNO1 vs. BL2×CNO2: n = 104, r = 0.60, p = 2.1 × 10^-11^; size change, BL×CNO1 vs. BL×CNO2: n = 64, r = 0.45, p = 1.8 × 10^-4^; linear correlations). Points represent CA1 place cells. **E)** Cumulative distribution function shows spatial correlation of rate maps from BL sessions on each recording day for place cells in hM3 (light blue) and Con (light orange) mice. Note that there was no difference between groups (spatial correlation, BL1×BL2: hM3 n = 126, median = 0.79, 95% CI, 0.69 – 0.82; Con n = 114, median = 0.79, 95% CI, 0.74 – 0.83; hM3 vs. Con, D* = 0.10, p = 0.57, two-sided Kolmogorov-Smirnov test), indicating that place cells in hM3 mice returned to their BL representations 12+ hrs. after CNO injection. For lower panels in (A) and (B), black dot: median; black bars: 95% confidence interval; filled curve: sampling-error distribution.

**Figure S5.**
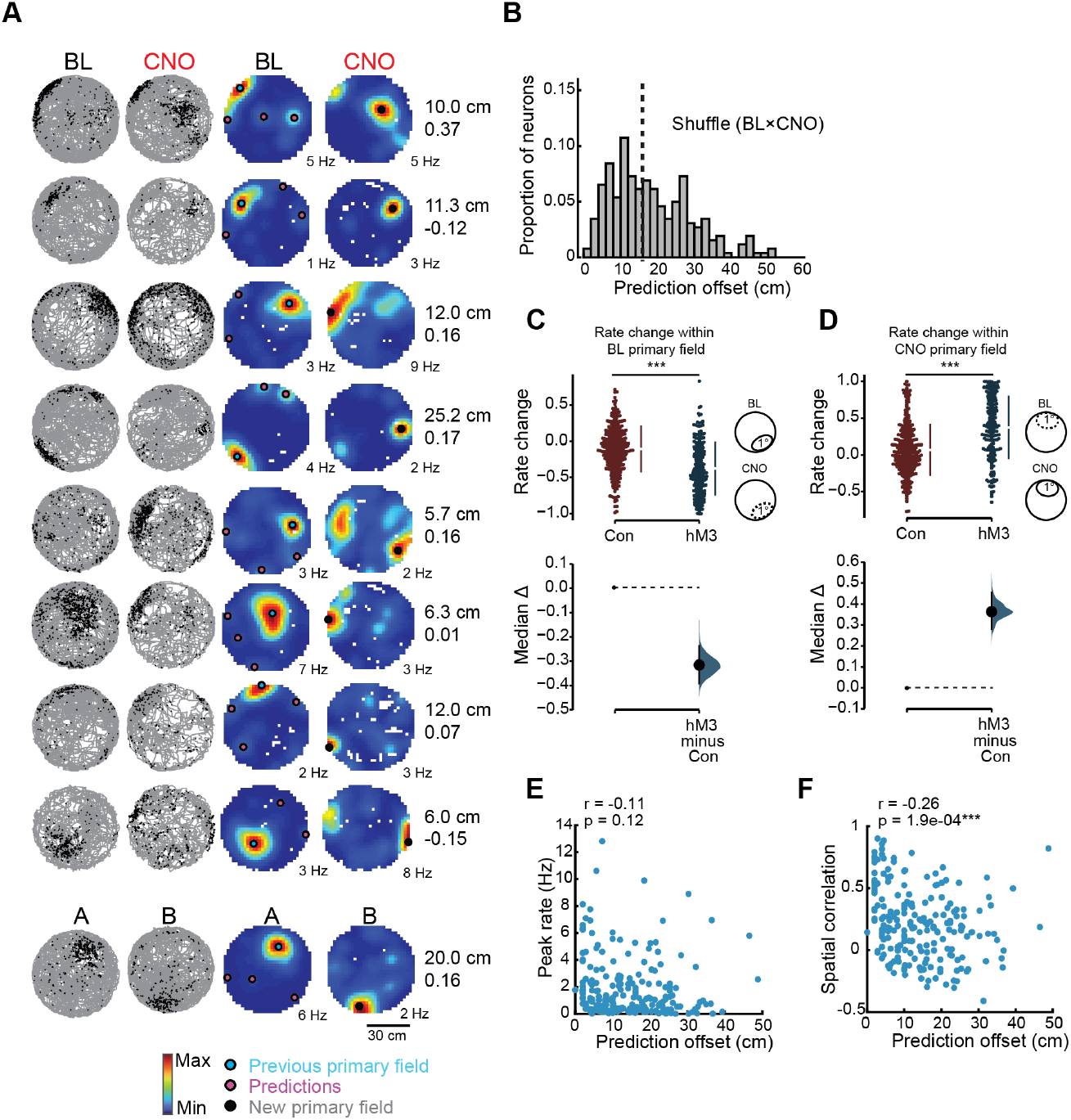
Predictable reorganization of place code despite robust artificial remapping. **A)** First eight rows show spike path plots and firing rate maps of BL and CNO sessions for eight CA1 place cells in hM3 mice (one cell per row). In spike path plots (first two columns), the mouse’s trajectory is shown in gray and action potentials are represented in black. For firing rate maps (last two columns), color indicates firing rate. Peak firing rate is noted below each rate map. Same convention for place field prediction as in Figure 3A (blue circles, primary place field in BL session; pink circles, predictions of place field location; black circles, primary place field in CNO session). The prediction offset and the spatial correlation between rate maps from the BL and CNO sessions are shown on the right for each cell. Bottom row shows spike path plots and firing rate maps for one CA1 place cell from a control mouse exposed to two distinct environments (A and B). Same convention as above. **B)** Histogram shows prediction offsets for a shuffled control dataset (see **Methods**). Note that prediction offsets were significantly lower in hM3 mice (see Figure 3C) than in the shuffled dataset (hM3 n = 204, median = 12.1 cm, 95% CI, 10.2 – 14.6 cm; shuffle n = 261, median = 16.1 cm, 95% CI, 14.4 – 17.9 cm; hM3 vs. shuffle, Z = 3.9, p = 5.6 × 10^-5^, one-sided Wilcoxon rank sum test). Vertical lines represent the median of the distribution. **C)** Panel shows the firing rate change within the primary place field from the BL session for place cells in hM3 (blue) and Con (red) mice. Between sessions, there was a significant decrease in firing rate within the BL primary field in hM3 mice relative to controls (Con n = 320, median = −0.10; hM3 n = 204, median = −0.42; Con vs. hM3, Z = 8.4, p = 5.3 × 10^-17^, two-sided Wilcoxon rank sum test). Points represent CA1 place cells; gapped lines represent mean ± standard deviation. Change refers to a difference score (see **Methods**). **D)** Panel shows the firing rate change within the primary place field from the CNO session for place cells in hM3 (blue) and Con (red) mice. Between sessions, there was a significant increase in firing rate within the CNO primary field in hM3 mice relative to controls (Con n = 320, median = 0.03; hM3 n = 204, median = 0.39; Con vs. hM3, Z = 8.4, p = 4.25 × 10^-17^, two-sided Wilcoxon rank sum test). Points represent CA1 place cells; gapped lines represent mean ± standard deviation. Change refers to a difference score (see **Methods**). **E)** Scatterplot shows no relationship between the peak firing rate in the predicted location in the BL session and the prediction offset for place cells in hM3 mice (n = 204, r = −0.11, p = 0.12, linear correlation). Points represent CA1 place cells. **F)** Scatterplot shows weak relationship between the degree of remapping following CNO injection and the prediction offset for place cells in hM3 mice (n = 204, r = −0.26, p = 1.9 × 10^-4^, linear correlation). Points represent CA1 place cells. *\*\*\*p* < 0.001. For lower panels in (C) and (D), black dot: median; black bars: 95% confidence interval; filled curve: sampling-error distribution.

**Figure S6.**
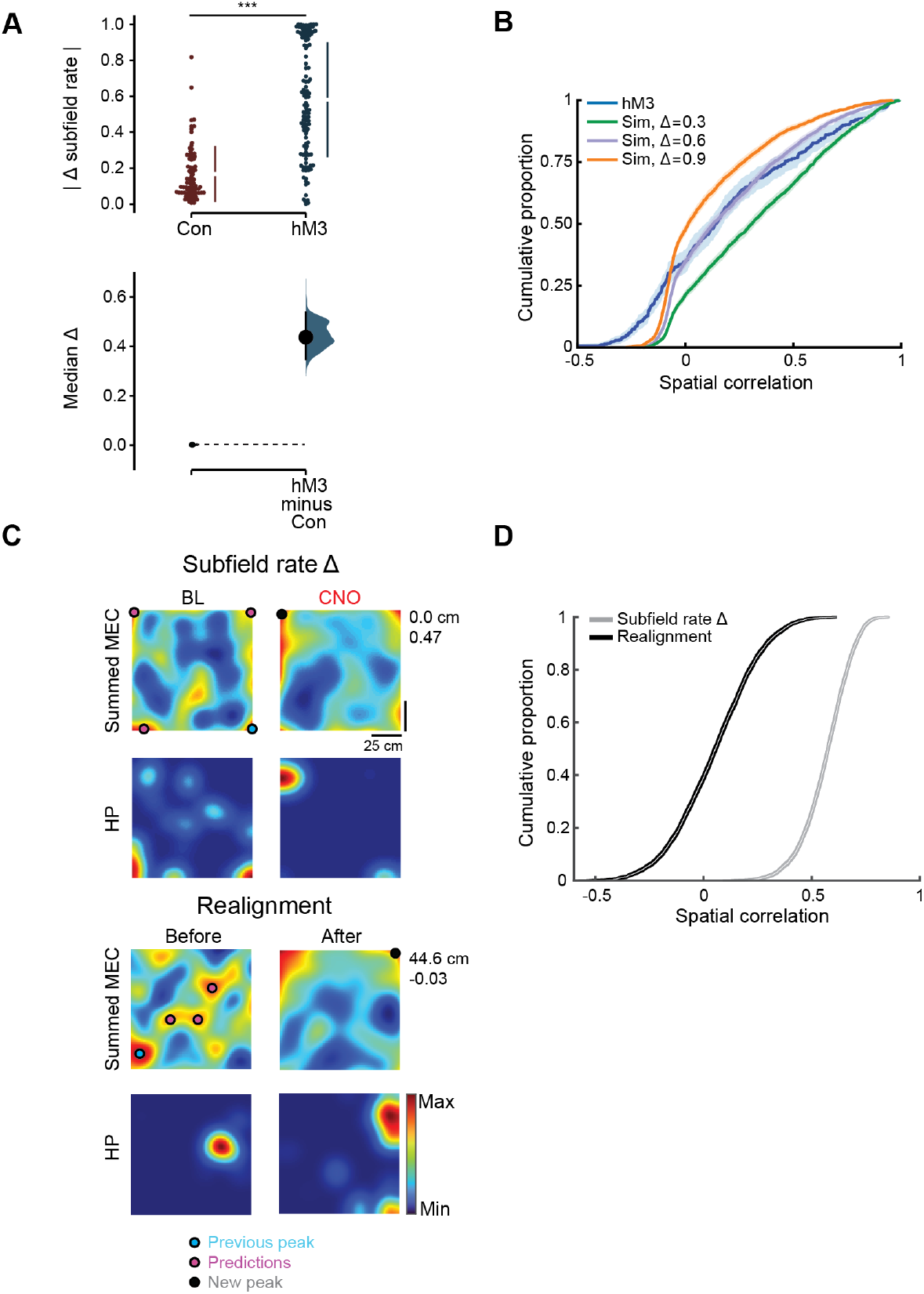
Grid subfield rate changes redistribute activity among spatially stable grid inputs. **A)** Panel shows distribution of grid subfield rate changes in hM3 (blue) and Con (red) mice between BL and CNO sessions. In our simulation of the grid-to-place cell transformation, grid subfield rates were modified by values drawn randomly from the distribution of grid field rate changes in hM3 mice (median = 0.55, 95% CI, 0.48 −0.65). Note that there was significantly more change in grid subfield rates between BL and CNO sessions in hM3 mice relative to controls (hM3 vs. Con: BL×CNO, hM3 n = 117, Con = 80, p = 2.7 × 10^-19^, two-sided Wilcoxon rank sum test). Top, points represent grid subfields; gapped lines represent mean ± standard deviation. Change refers to an absolute difference score (see **Methods**). Bottom, test statistic is the median difference, shown on the y axis as a bootstrap sampling distribution. Black dot: median; black bars: 95% confidence interval; filled curve: sampling-error distribution. p value is from Wilcoxon rank sum test; *\*\*\*p* < 0.001. **B)**Cumulative distribution function shows spatial correlation between rate maps from BL and CNO sessions for place cells in hM3 mice (blue) and simulated place cells. When grid subfield rates were modified by values drawn randomly from the distribution of grid field rate changes in hM3 mice (Sim Δ = 0.6, purple), the extent of remapping was similar for simulated place cells and place cells in hM3 mice (spatial correlation, BL×CNO: hM3 n = 394, median = 0.14, 95% CI, 0.09 – 0.17; Sim Δ = 0.6 n = 1,975, median = 0.13, 95% CI, 0.11 – 0.16; hM3 vs. Sim Δ = 0.6, Z = 1.7, p = 0.09, two-sided Wilcoxon rank sum test). Decreasing (Sim Δ = 0.3, green) or increasing (Sim Δ = 0.9, orange) the extent of grid subfield rate changes (by adjusting the median of the distribution from which subfield rates were sampled) modulated the degree of remapping among simulated place cells (spatial correlation, BL×CNO: Sim Δ = 0.9 n = 2,033, median = 0.01, 95% CI, 0.00 – 0.03; Sim Δ = 0.3 n = 1,897, median = 0.31, 95% CI, 0.28 – 0.33). **C)** Panels show excitation maps representing the summed grid cell input to a single simulated place cell before and after grid subfield rate change (top) or independent realignment of grid modules (bottom). Excitation maps (top row) depict strength of summed grid input (from blue to red). Color in corresponding place cell rate maps (bottom row) indicates firing rate. To predict changes in the location of summed grid inputs (rather than hippocampal place fields), we used the same method as in Figure 3A (blue circles, previous peak; pink circles, predictions; black circles, new peak). The prediction offset and the spatial correlation between excitation maps from each session are shown on the right. Note that grid subfield rate changes typically caused the location of the primary field to shift to an alternate peak in the input pattern rather than a random location, resulting in low prediction offsets and high spatial correlations between sessions (median prediction offset = 16.0 cm, 95% CI, 13.0 – 19.0 cm). The location of the primary field typically shifted to an unpredicted location following independent realignment of grid modules, resulting in high prediction offsets and low spatial correlations between sessions (median prediction offset = 40.2 cm, 95% CI, 39.0 – 41.2 cm). The prediction offset was significantly lower after grid subfield rate changes than independent realignment (subfield rate change n = 3,730, independent realignment n = 4,616, Z = 21.8, p = 9.3 × 10^-106^, two-sided Wilcoxon rank sum test). **D)** Cumulative distribution functions show spatial correlation between excitation maps before and after grid subfield rate change (gray) or independent realignment of grid modules (black). Grid subfield rate changes resulted in a predictable reorganization of grid cell input, resulting in significantly higher spatial correlation between excitation maps from each session than following independent realignment (spatial correlation: subfield rate change n = 5,000, median = 0.575, 95% CI, 0.571 – 0.579; independent realignment n = 5,000, median = 0.054, 95% CI, 0.047 – 0.061; subfield rate change vs. independent realignment, D* = 0.91, p < 2.2 × 10^-16^, two-sided Kolmogorov-Smirnov test).

**Figure S7.**
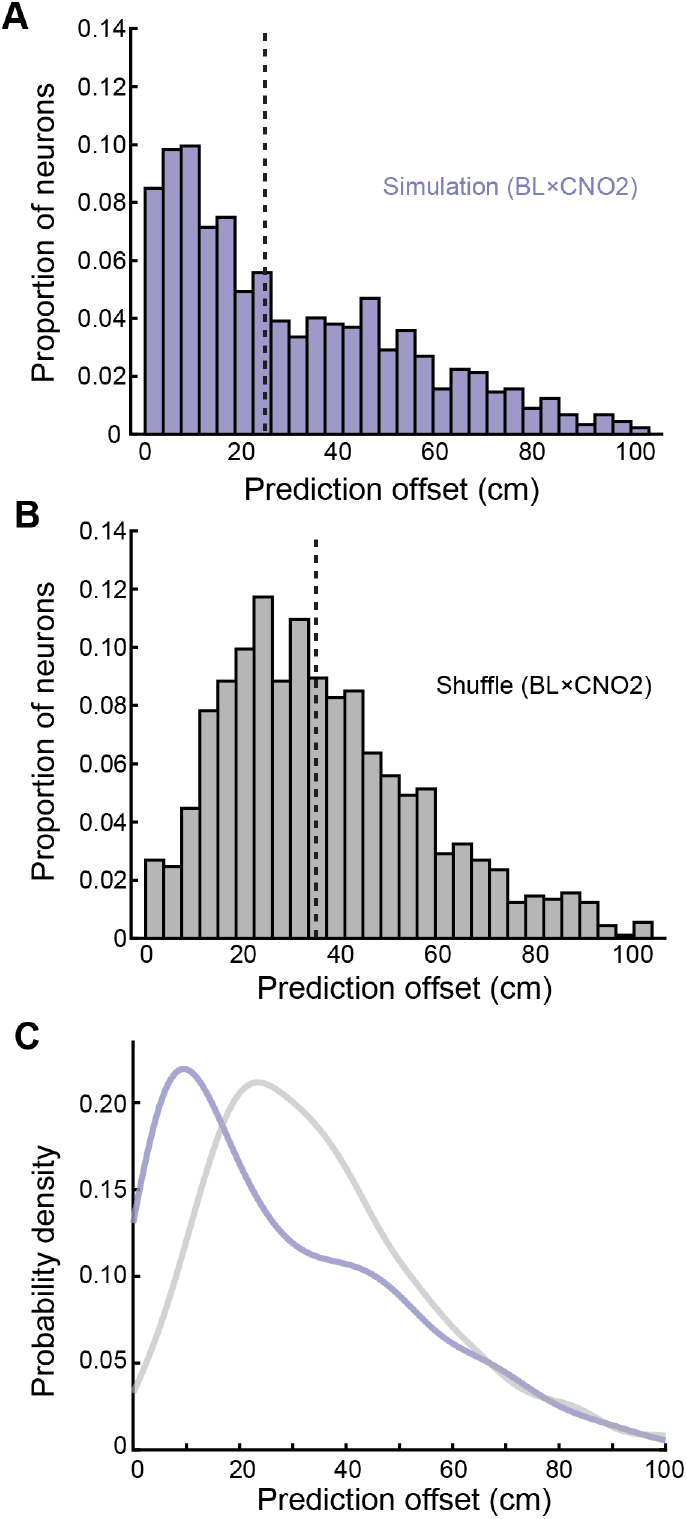
Grid subfield rate changes elicit predictable and reproducible reorganization of hippocampal place fields. **A)** Histogram shows prediction offsets between BL and CNO2 for simulated place cells. Note that we were able to predict place field locations equally well during both runs of the simulation (prediction offset: BL×CNO2 n = 895, median = 22.2 cm, 95% CI, 19.4 – 24.1 cm; BL×CNO1 vs. BL×CNO2, Z = 1.0, p = 0.31, two-sided Wilcoxon rank sum test). Vertical line represents the median of the distribution. **B)** Histograms show prediction offsets between BL and CNO2 for a shuffled control dataset (see **Methods**; BL×CNO2 n = 1,218, median = 31.7 cm, 95% CI, 30.2 – 33.1 cm). Note that prediction offsets for simulated place cells between BL and CNO2 were significantly lower for the shuffled control dataset (simulation vs. shuffle, Z = 8.7, p = 4.4 × 10^-18^, two-sided Wilcoxon rank sum test). Vertical line represents the median of the distribution. **C)** Kernel smoothed density estimate of prediction offset for simulated place cells (purple) and a shuffled control dataset (gray) between the BL session and CNO2 (BL×CNO2).

## Notes

### Competing Interest Statement

The authors have declared no competing interest.

## References

Agmon, H. and Burak, Y. (2020). A theory of joint attractor dynamics in the hippocampus and the entorhinal cortex accounts for artificial remapping and grid cell field-to-field variability. Elife, 9. doi: 10.7554/eLife.56894.

Alexander, G. M., Rogan, S. C., Abbas, A. I., Armbruster, B. N., Pei, Y., Allen, J. A., Nonneman, R. J., Hartmann, J., Moy, S. S., Nicolelis, M. A., McNamara, J. O., and Roth, B. L. (2009). Remote control of neuronal activity in transgenic mice expressing evolved g protein-coupled receptors. Neuron, 63(1):27–39. doi: 10.1016/j.neuron.2009.06.014.

Alme, C. B., Miao, C., Jezek, K., Treves, A., Moser, E. I., and Moser, M. B. (2014). Place cells in the hippocampus: Eleven maps for eleven rooms. Proc Natl Acad Sci U S A, 111(52):18428–18435. doi: 10.1073/pnas.1421056111.

Anderson, M. I. and Jeffery, K. J. (2003). Heterogeneous modulation of place cell firing by changes in context. J Neurosci, 23 (26):8827–35. doi: 10.1523/JNEUROSCI.23-26-08827.2003.

Bittner, K. C., Grienberger, C., Vaidya, S. P., Milstein, A. D., Macklin, J. J., Suh, J., Tonegawa, S., and Magee, J. C. (2015). Conjunctive input processing drives feature selectivity in hippocampal ca1 neurons. Nat Neurosci, 18(8):1133–42. doi: 10.1038/nn.4062.

Bittner, K. C., Milstein, A. D., Grienberger, C., Romani, S., and Magee, J. C. (2017). Behavioral time scale synaptic plasticity underlies ca1 place fields. Science, 357(6355):1033–1036. doi: 10.1126/science.aan3846.

Bostock, E., Muller, R. U., and Kubie, J. L. (1991). Experience-dependent modifications of hippocampal place cell firing. Hippocampus, 1(2):193–205. doi: 10.1002/hipo.450010207.

Brandon, M. P., Koenig, J., Leutgeb, J. K., and Leutgeb, S. (2014). New and distinct hippocampal place codes are generated in a new environment during septal inactivation. Neuron, 82(4):789–796. doi: 10.1016/j.neuron.2014.04.013.

Butler, W. N., Hardcastle, K., and Giocomo, L. M. (2019). Remembered reward locations restructure entorhinal spatial maps. Science, 363(6434):1447–1452. doi: 10.1126/science.aav5297.

Cappaert, N. L. M., Van Strien, N. M., and Witter, M. P. Chapter 20 - hippocampal formation. In Paxinos, G., editor, The Rat Nervous System, pages 511–573. Academic Press, fourth edition edition, (2015). doi: 10.1016/B978-0-12-374245-2.00020-6.

Cohen, J. D., Bolstad, M., and Lee, A. K. (2017). Experience-dependent shaping of hippocampal ca1 intracellular activity in novel and familiar environments. eLife, 6:e23040. doi: 10.7554/eLife.23040.

de Almeida, L., Idiart, M., and Lisman, J. E. (2009). The input-output transformation of the hippocampal granule cells: from grid cells to place fields. J Neurosci, 29(23):7504–7512. doi: 10.1523/jneurosci.6048-08.2009.

de Almeida, L., Idiart, M., and Lisman, J. E. (2012). The single place fields of CA3 cells: a two-stage transformation from grid cells. Hippocampus, 22(2):200–208. doi: 10.1002/hipo.20882.

Diamantaki, M., Coletta, S., Nasr, K., Zeraati, R., Laturnus, S., Berens, P., Preston-Ferrer, P., and Burgalossi, A. (2018). Manipulating hippocampal place cell activity by single-cell stimulation in freely moving mice. Cell Rep, 23(1):32–38. doi: 10.1016/j.celrep.2018.03.031.

Diehl, G., Hon, O., Leutgeb, S., and Leutgeb, J. (2017). Grid and nongrid cells in medial entorhinal cortex represent spatial location and environmental features with complementary coding schemes. Neuron, 94:83–92. doi: 10.1016/j.neuron.2017.03.004.

Dunn, B., Wennberg, D., Huang, Z., and Roudi, Y. (2017). Grid cells show field-to-field variability and this explains the aperiodic response of inhibitory interneurons. bioRxiv, 101899.

Eichenbaum, H. (2017). The role of the hippocampus in navigation is memory. Journal of Neurophysiology, 117(4):1785–1796. doi: 10.1152/jn.00005.2017.

Eichenbaum, H., Dudchenko, P., Wood, E., Shapiro, M., and Tanila, H. (1999). The hippocampus, memory, and place cells: is it spatial memory or a memory space? Neuron, 23(2):209–26.

Fuhs, M. C. and Touretzky, D. S. (2006). A spin glass model of path integration in rat medial entorhinal cortex. J Neurosci, 26 (16):4266–4276. doi: 10.1523/JNEUROSCI.4353-05.2006.

Fyhn, M., Hafting, T., Treves, A., Moser, M. B., and Moser, E. I. (2007). Hippocampal remapping and grid realignment in entorhinal cortex. Nature, 446(7132):190–194. doi: 10.1038/nature05601.

Gray, C. M., Maldonado, P. E., Wilson, M., and McNaughton, B. (1995). Tetrodes markedly improve the reliability and yield of multiple single-unit isolation from multi-unit recordings in cat striate cortex. J Neurosci Methods, 63(1-2):43–54. doi: 10.1016/0165-0270(95)00085-2.

Hafting, T., Fyhn, M., Molden, S., Moser, M. B., and Moser, E. I. (2005). Microstructure of a spatial map in the entorhinal cortex. Nature, 436(7052):801–806. doi: 10.1038/nature03721.

Ho, J., Tumkaya, T., Aryal, S., Choi, H., and Claridge-Chang, A. (2019). Moving beyond p values: data analysis with estimation graphics. Nat Methods, 16(7):565–566. doi: 10.1038/s41592-019-0470-3.

Ismakov, R., Barak, O., Jeffery, K., and Derdikman, D. (2017). Grid cells encode local positional information. Curr Biol, 27: 2337–2343. doi: 10.1016/j.cub.2017.06.034.

Kanter, B. R., Lykken, C. M., Avesar, D., Weible, A., Dickinson, J., Dunn, B., Borgesius, N. Z., Roudi, Y., and Kentros, C. G. (2017). A novel mechanism for the grid-to-place cell transformation revealed by transgenic depolarization of medial entorhinal cortex layer II. Neuron, 93(6):1480–1492. doi: 10.1016/j.neuron.2017.03.001.

Kentros, C. (2006). Hippocampal place cells: the “where” of episodic memory? Hippocampus, 16(9):743–54. doi: 10.1002/hipo.20199.

Kentros, C. G., Agnihotri, N. T., Streater, S., Hawkins, R. D., and Kandel, E. R. (2004). Increased attention to spatial context increases both place field stability and spatial memory. Neuron, 42(2):283–295.

Knierim, J. J. (2002). Dynamic interactions between local surface cues, distal landmarks, and intrinsic circuitry in hippocampal place cells. The Journal of Neuroscience, 22(14):6254–6264. doi: 10.1523/JNEUROSCI.22-14-06254.2002.

Langston, R. F., Ainge, J. A., Couey, J. J., Canto, C. B., Bjerknes, T. L., Witter, M. P., Moser, E. I., and Moser, M. B. (2010). Development of the spatial representation system in the rat. Science, 328(5985):1576–1580. doi: 10.1126/science.1188210.

Lee, D., Lin, B. J., and Lee, A. K. (2012). Hippocampal place fields emerge upon single-cell manipulation of excitability during behavior. Science, 337(6096):849–853. doi: 10.1126/science.1221489.

Leutgeb, S., Leutgeb, J. K., Barnes, C. A., Moser, E. I., McNaughton, B. L., and Moser, M. B. (2005). Independent codes for spatial and episodic memory in hippocampal neuronal ensembles. Science, 309(5734):619–623. doi: 10.1126/science.1114037.

Liao, Z., Gonzalez, K. C., Li, D. M., Yang, C. M., Holder, D., McClain, N. E., Zhang, G., Evans, S. W., Chavarha, M., Simko, J., Makinson, C. D., Lin, M. Z., Losonczy, A., and Negrean, A. (2024). Functional architecture of intracellular oscillations in hippocampal dendrites. Nature Communications, 15(1):6295. doi: 10.1038/s41467-024-50546-z.

Lykken, C. M., Kanter, B. R., Nagelhus, A., Carpenter, J., Guardamagna, M., Moser, E. I., and Moser, M.-B. (2025). Functional independence of entorhinal grid cell modules enables remapping in hippocampal place cells. bioRxiv, page 2025.09.24.677985. doi: 10.1101/2025.09.24.677985.

Lyttle, D., Gereke, B., Lin, K. K., and Fellous, J. M. (2013). Spatial scale and place field stability in a grid-to-place cell model of the dorsoventral axis of the hippocampus. Hippocampus, 23(8):729–744. doi: 10.1002/hipo.22132.

McKenzie, S., Huszár, R., English, D. F., Kim, K., Christensen, F., Yoon, E., and Buzsáki, G. (2021). Preexisting hippocampal network dynamics constrain optogenetically induced place fields. Neuron. doi: 10.1016/j.neuron.2021.01.011.

McNaughton, B. L., Battaglia, F. P., Jensen, O., Moser, E. I., and Moser, M.-B. (2006). Path integration and the neural basis of the ‘cognitive map’. Nat Rev Neurosci, 7(8):663–678.

Monaco, J. D. and Abbott, L. F. (2011). Modular realignment of entorhinal grid cell activity as a basis for hippocampal remapping. J Neurosci, 31(25):9414–9425. doi: 10.1523/jneurosci.1433-11.2011.

Muller, R. U. and Kubie, J. L. (1987). The effects of changes in the environment on the spatial firing of hippocampal complex-spike cells. J Neurosci, 7(7):1951–1968.

O’Keefe, J. and Dostrovsky, J. (1971). The hippocampus as a spatial map. preliminary evidence from unit activity in the freely-moving rat. Brain Res, 34(1):171–175.

O’Keefe, J. and Nadel, L. The Hippocampus as a Cognitive Map. Clarendon Press, Oxford, (1978). ISBN 0-19-857206-9.

Ormond, J. and McNaughton, B. L. (2015). Place field expansion after focal mec inactivations is consistent with loss of fourier components and path integrator gain reduction. Proc Natl Acad Sci U S A, 112(13):4116–4121. doi: 10.1073/pnas.1421963112.

Ormond, J., Serka, S. A., and Johansen, J. P. (2023). Enhanced reactivation of remapping place cells during aversive learning. J Neurosci, 43(12):2153–2167. doi: 10.1523/JNEUROSCI.1450-22.2022.

Redman, W. T., Acosta-Mendoza, S., Wei, X.-X., and Goard, M. J. (2025). Robust variability of grid cell properties within individual grid modules enhances encoding of local space. eLife, 13:RP100652. doi: 10.7554/eLife.100652.

Robinson, N. T. M., Descamps, L. A. L., Russell, L. E., Buchholz, M. O., Bicknell, B. A., Antonov, G. K., Lau, J. Y. N., Nutbrown, R., Schmidt-Hieber, C., and Hausser, M. (2020). Targeted activation of hippocampal place cells drives memory-guided spatial behavior. Cell, 183(6):1586–1599 e10. doi: 10.1016/j.cell.2020.09.061.

Rolls, E. T., Stringer, S. M., and Elliot, T. (2006). Entorhinal cortex grid cells can map to hippocampal place cells by competitive learning. Network, 17(4):447–465. doi: 10.1080/09548980601064846.

Sargolini, F., Fyhn, M., Hafting, T., McNaughton, B. L., Witter, M. P., Moser, M. B., and Moser, E. I. (2006). Conjunctive representation of position, direction, and velocity in entorhinal cortex. Science, 312(5774):758–762. doi: 10.1126/science.1125572.

Savelli, F. and Knierim, J. J. (2010). Hebbian analysis of the transformation of medial entorhinal grid-cell inputs to hippocampal place fields. J Neurophysiol, 103(6):3167–3183. doi: 10.1152/jn.00932.2009.

Savelli, F., Yoganarasimha, D., and Knierim, J. J. (2008). Influence of boundary removal on the spatial representations of the medial entorhinal cortex. Hippocampus, 18(12):1270–1282. doi: 10.1002/hipo.20511.

Schlesiger, M. I., Boublil, B. L., Hales, J. B., Leutgeb, J. K., and Leutgeb, S. (2018). Hippocampal global remapping can occur without input from the medial entorhinal cortex. Cell Rep, 22(12):3152–3159. doi: 10.1016/j.celrep.2018.02.082.

Scoville, W. B. and Milner, B. (1957). Loss of recent memory after bilateral hippocampal lesions. J Neurol Neurosurg Psychiatry, 20(1):11–21.

Shapiro, e. a. M. (1997). Cues that hippocampal place cells encode: Dynamic and hierarchical representation of local and distal stimuli. Hippocampus, 7(6):624–42.

Solstad, T., Moser, E. I., and Einevoll, G. T. (2006). From grid cells to place cells: a mathematical model. Hippocampus, 16(12): 1026–1031. doi: 10.1002/hipo.20244.

Solstad, T., Boccara, C. N., Kropff, E., Moser, M. B., and Moser, E. I. (2008). Representation of geometric borders in the entorhinal cortex. Science, 322(5909):1865–1868. doi: 10.1126/science.1166466.

Tanila, e. a. H. (1997). Discordance of spatial representation in ensembles of hippocampal place cells. Hippocampus, 7(6): 613–623.

Taube, J. S., Muller, R. U., and Ranck, J., J. B. (1990). Head-direction cells recorded from the postsubiculum in freely moving rats. i. description and quantitative analysis. J Neurosci, 10(2):420–35. doi: 10.1523/JNEUROSCI.10-02-00420.1990.

Treves, A. and Rolls, E. T. (1991). What determines the capacity of autoassociative memories in the brain? Network: Computation in Neural Systems, 2(4):371–397. doi: 10.1088/0954-898X_2_4_004.

Treves, S.. (2009). The role of competitive learning in the generation of dg fields from ec inputs. Cogn Neurodyn.

Yasuda, M. and Mayford, M. R. (2006). CaMKII activation in the entorhinal cortex disrupts previously encoded spatial memory. Neuron, 50(2):309–318. doi: 10.1016/j.neuron.2006.03.035.

